# Next-generation anti-PD-L1/IL-15 immunocytokine elicits superior antitumor immunity in cold tumors with minimal toxicity

**DOI:** 10.1101/2023.08.02.551593

**Authors:** Wenqiang Shi, Nan Liu, Zexin Liu, Yuqi Yang, Qiongya Zeng, Yang Wang, Luyao Song, Jianwei Zhu, Huili Lu

## Abstract

Immunocytokines, such as anti-PD-L1/IL-15, have shown promising efficacy in preclinical studies, but their clinical development still faces severe safety concerns, with the problem not easily overcome by simply reducing the cytokine activity. We proposed a next-generation immunocytokine concept of designing a tumor-conditional anti-PD-L1/IL-15 prodrug (LH05), which innovatively masks IL-15 with steric hindrance of its flanking moieties of anti-PD-L1 and IL-15Rα-sushi domain. The design successfully attenuated the ‘cytokine sink’ effect of IL-15 and resulted in a significantly reduced systemic toxicity when compared to wild-type anti-PD-L1/IL-15. LH05 would be specifically cleaved in the tumor microenvironment (TME) to release the active IL-15/IL-15Rα-sushi domain (ILR) in a proteolytic cleavage-dependent manner and exhibited potent antitumor effects in mouse syngeneic models. Mechanistically, the antitumor efficacy of LH05 was dependent on both innate and adaptive immunity, which altered the TME to Th1-type by recruiting and stimulating both NK and CD8^+^ T cells and fired up cold tumors. LH05 also showed superior efficacy in restoring immunotherapy response in a refractory U251 xenograft model. Collectively, we introduced a novel next-generation immunocytokine strategy for tumor immunotherapy, contributing to the establishment of optimal treatment for patients with resistance to immune checkpoint inhibitors or cold tumors.

## Introduction

Many cytokines have demonstrated potent antitumor activity in preclinical studies, but their clinical utility is limited due to their short half-lives and systemic toxicity [1]. Antibody–cytokine fusion proteins (immunocytokines) delivering these immunostimulatory payloads to tumor lesions can substantially broaden the therapeutic window of cytokine therapy. Additionally, combining antibody and cytokine can generate synergistic antitumor effects [2]. Some immunocytokines based on IL-2, IL-12, TNF-α, etc. have been investigated in clinical trials, among which EDB (fibronectin extradomain)-specific immunocytokines with TNF-α or IL-2 payloads have progressed to phase III trials (NCT02938299 and NCT03567889) [3]. The antibodies of previously published immunocytokines mostly targeted highly expressed targets in the tumor microenvironment (TME), such as fibronectin and fibroblast activation protein [4]. With the considerable advancements of immune checkpoint inhibitors (ICIs) in cancer immunotherapy, antibodies targeting immune checkpoints have recently emerged as the main protagonists of immunocytokines [5,6].

However, immunocytokines can be trapped by cognate receptors in circulation before reaching their target cells (so-called “sink effect”) [7]. This off-target effect of immunocytokines can lead to systemic toxicity. Moreover, only a small fraction of the immunocytokine can be taken up by the neoplastic lesion (in the best cases, 0.01%– 0.1% injected dose/g of tumor), often resulting in toxicity profiles similar to that of parental cytokine [8]. Notably, patients treated with KD033 (a PD-L1/IL-15 bispecific molecule) at a dose of 50 μg/kg experienced severe lymphocytopenia, despite the fact that this dose is much lower than the clinical dose of anti-PD-L1 (10–20 mg/kg) [9,10]. It is crucial to develop novel strategies to overcome safety challenges of immunocytokines and promote their clinical applications.

To reduce systemic toxicity of immunocytokines, cytokines should be engineered to reduce affinity for their cognate receptors. For example, AcTaferon, comprising human IFNα2 (Q124R) that is 100 folds less active on mouse cells as compared to murine IFNα fused to anti-CD20, demonstrated a strong antitumor activity without any associated toxicity, in contrast with wild-type IFNα2. Additionally, the prodrug strategy that can selectively release cytokine activity in the TME represents a promising approach for the development of next-generation immunocytokine. Spatial hindrance and affinity peptides are among the most popular masking strategies for biomolecules, including antibodies or cytokines [11,12]. Using a cleavable linker, Fu et al. engineered masked IL-2, IL-12, IL-15, and type I IFN with their natural receptors [13–16]. These pro-cytokines reactivate after being cleaved by tumor-associated enzymes within the TME. Although the receptor-masked strategy can reduce the peripheral activity of the cytokine, the introduction of the masking receptor complicates the structure. Moreover, the cleaved receptors may have unfavorable influence on sufficiently restoring the cytokine activity, since cytokines and their receptors have a high affinity for each other [17].

IL-15 is a highly attractive immunostimulatory cytokine due to its remarkable activity in treating various cancer types [18, 19]. Immunocytokines with IL-15 as a payload have shown great prospect in clinical applications, including KD033 and BJ-001, which fuse IL-15 with anti-PD-L1 antibody and integrin-targeting RGD peptide, respectively [20, 21]. We have previously developed an anti-PD-L1/IL-15 immunocytokine (LH01), which can overcome anti-PD-L1 resistance and elicit both innate and adaptive immune responses. However, LH01 also induces systemic toxicity similar to that of IL-15 [22].

To improve the drug-like properties of immunocytokines and treat cold tumors resistant to existing immunotherapy regimens, in the present study, we designed next-generation anti-PD-L1/IL-15 (LH05), a prodrug masking IL-15, with an innovative steric hindrance strategy. This prodrug can be preferentially cleaved within the TME by a tumor-associated protease to release the reactivated IL-15/IL-15Rα-sushi domain (ILR, an IL-15 superagonist) [23], which would have more pleiotropic anticancer effects as compared to being bound to the antibody. Using this strategy, LH05 addressed the safety concerns of IL-15-based immunocytokines and enhanced its efficacy, providing a preclinical proof of concept for the development of next-generation immunocytokines.

## Results

### Systemic toxicity restricts the efficacy of anti-PD-L1/IL-15 immunocytokine in cold tumor

We have demonstrated the potent antitumor efficacy of anti-PD-L1/IL-15 immnocytokine (LH01) in syngeneic murine tumor and xenograft models in previous studies. In the present study, we further studied the therapeutic effect of LH01 in treating cold tumors. We observed that LH01 has a dose-related therapeutic outcome and toxicity in RM-1 syngeneic prostate model with a “cold” immune landscape. LH01 was well tolerated at 2.5 mg/kg but it only exerted a slight antitumor activity (Fig. 1A-C). When the dosage was increased to 5 mg/kg, LH01 demonstrated a significant antitumor activity (Fig. 1A). However, it induced significant body weight loss and even death (half of the mice died) after two treatments (Fig. 1B and C). In short, the dose-limiting toxicities hinder the therapeutic efficacy of LH01 in treating cold tumors.

**Fig. 1.**
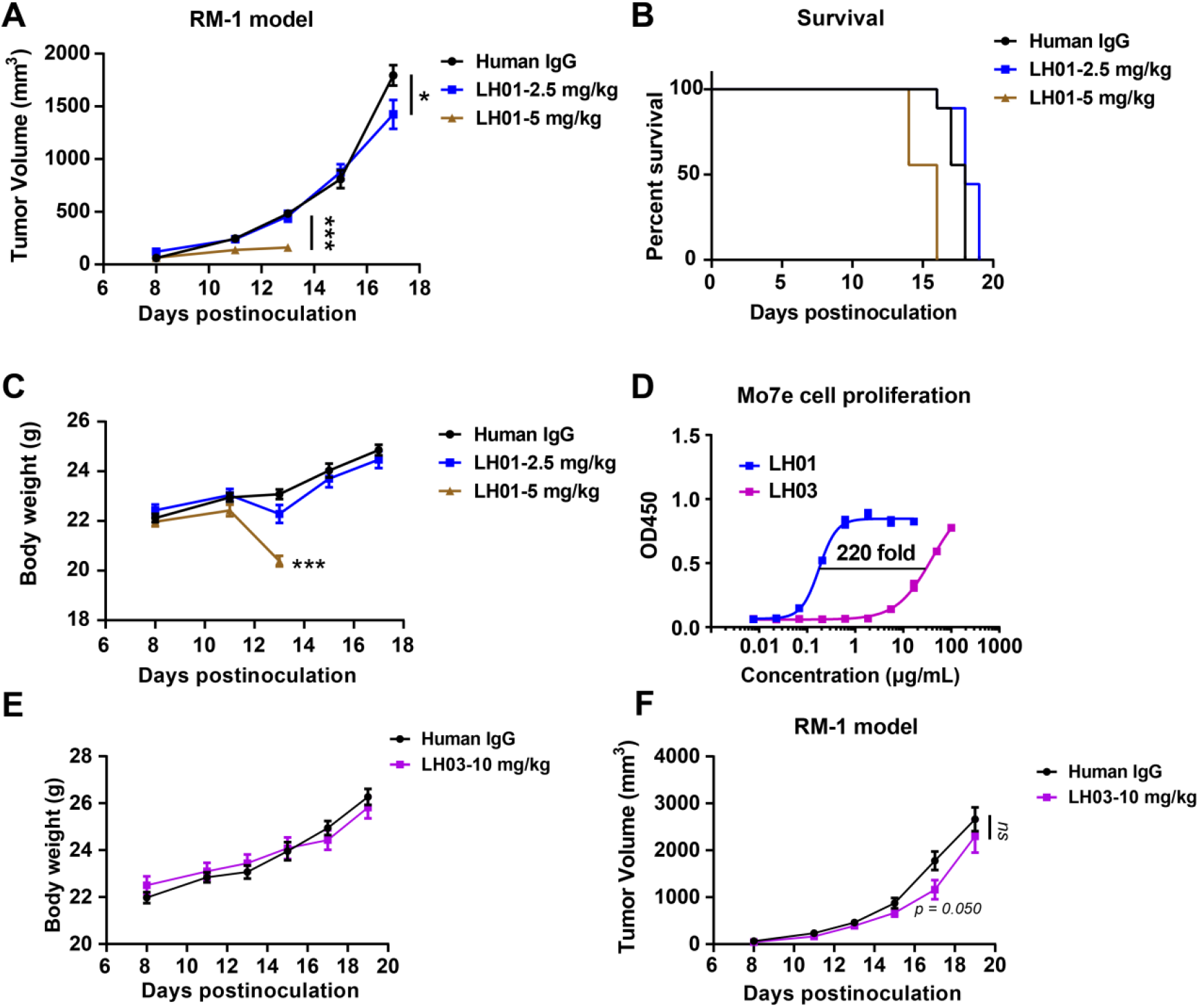
The limited antitumor efficacy of LH01 and LH03 in RM-1 cold tumor. **(A-C)** RM-1 tumor cells (5 × 10^5^) were subcutaneously implanted into the right flank of male C57BL/6J mice. Mice were randomized into three groups based on tumor size and treatment initiated when tumors reached 50-100 mm^3^ (n = 9). Mice were intravenously injected with IgG control (10 mg/kg) or LH01 (2.5 mg/kg or 5 mg/kg) on days 9, 12, and 15. (A) Tumor growth curves were plotted over time. Mouse survival (B) and body weight (C) were monitored. **(D)** The proliferative potential of LH03 was compared with LH01 in human Mo7e cells. Data were analyzed using the four parameter fit logistic equation to calculate the EC_50_ values. **(E-F)** Male C57BL/6J mice were inoculated with 5 × 10^5^ RM-1 tumor cells. When tumors reached 50-100 mm^3^, mice were intravenously injected with IgG control (10 mg/kg) or LH03 (10 mg/kg) on days 9, 12, and 15. (E) Tumor progression curves and body weight (F) were depicted. All graphs show the mean ±SEM. *p < 0.05; **p < 0.01; ***p < 0.001; ns, not significant.

**Table 1.** Pharmacokinetic parameters of LH01 and LH05.

| Parameters | LH01 | LH05 |
| --- | --- | --- |
| Half-life (T1/2), h | 3.45 | 8.40 |
| Cmax, ng/mL | 7734.76 | 8237.93 |
| AUC (0→ ∞), ng×h/mL | 28364.2 | 48535.74 |
| MRT, h | 4.57 | 10.41 |

\* Calculated with PK Solver 2.0 for a non-compartmental model.
\* C<sub>max</sub>, peak concentration; AUC, area under the curve; MRT, mean resident time.

To improve the efficacy and avoid toxicity, we first attempted to mitigate the IL-15 activity. The lower biological activity allows the use of higher doses, which may provide an avenue to create in *vivo* selectivity. We then engineered an anti-PD-L1 fusion (LH03), wherein IL-15 is fused to the C-terminus of anti-PD-L1 and the N-terminus of sushi domain via an engineered linker. LH03 demonstrated decreased affinity toward the IL-15Rβ as compared with LH01 (Supplementary Fig. 1). Besides, LH03 (EC_50_ = 39.24 μg/mL or 194.2 nM) induced 220-fold less proliferative activity than LH01 (EC_50_ = 0.177 μg/mL or 0.88 nM) in human Mo7e cells, suggesting successfully masked IL-15 immuno-stimulatory activity (Fig. 1D). As expected, the safety was largely improved and no body weight loss was observed for the LH03 group even at a dose of 10 mg/kg (Fig. 1E). However, there was also no therapeutic efficacy was observed, and the tumor growth was close to that of the control (Fig. 1F). Additionally, LH03 at 10 mg/kg exerts no significant antitumor effects but it has good tolerability in the MC38 and Renca models (Supplementary Fig. 2). Altogether, our findings demonstrated that balancing the toxicity and efficacy of immunocytokines by reducing cytokine activity is difficult. A novel strategy or design is necessary to address the challenges of immunocytokine drug development.

### Steric masking of IL-15 activity and in vitro activation of LH05 by tumor-specific protease

We sought to develop an engineered IL-15 blockade that retains its antitumor activity while limiting systemic exposure. Considering that ILR complex was reported as an IL-15 superagonist, we devised a next-generation IL-15-based immunocytokine (LH05) by incorporating a protease-cleavable linker between the antibody and ILR, which can mask IL-15 activity by steric hindrance caused by the Fc fragment and the sushi domain. The cleavable linker was chosen for its protease sensitivity, which is overexpressed in various human carcinomas: urokinase-type plasminogen activator (uPA) (Supplementary Fig. 3). It would act as a switch for IL-15 activity. Before its cleavage, IL-15 is shielded by the joint forces of Fc and sushi domain. After its cleavage, ILR would be released, restoring the antitumor activity. The proposed mechanism of action of LH05 is illustrated in Fig. 2A. We simulated the conformational structures of LH01 and LH05 by using AlphaFold, which showed that the IL-15 portion in LH01 was free and the receptor-binding sites were exposed. Contrarily, the IL-15 portion of LH05 was restricted due to steric hindrance caused by the Fc fragment and the sushi domain (Supplementary Fig. 4A and B).

**Fig. 2.**
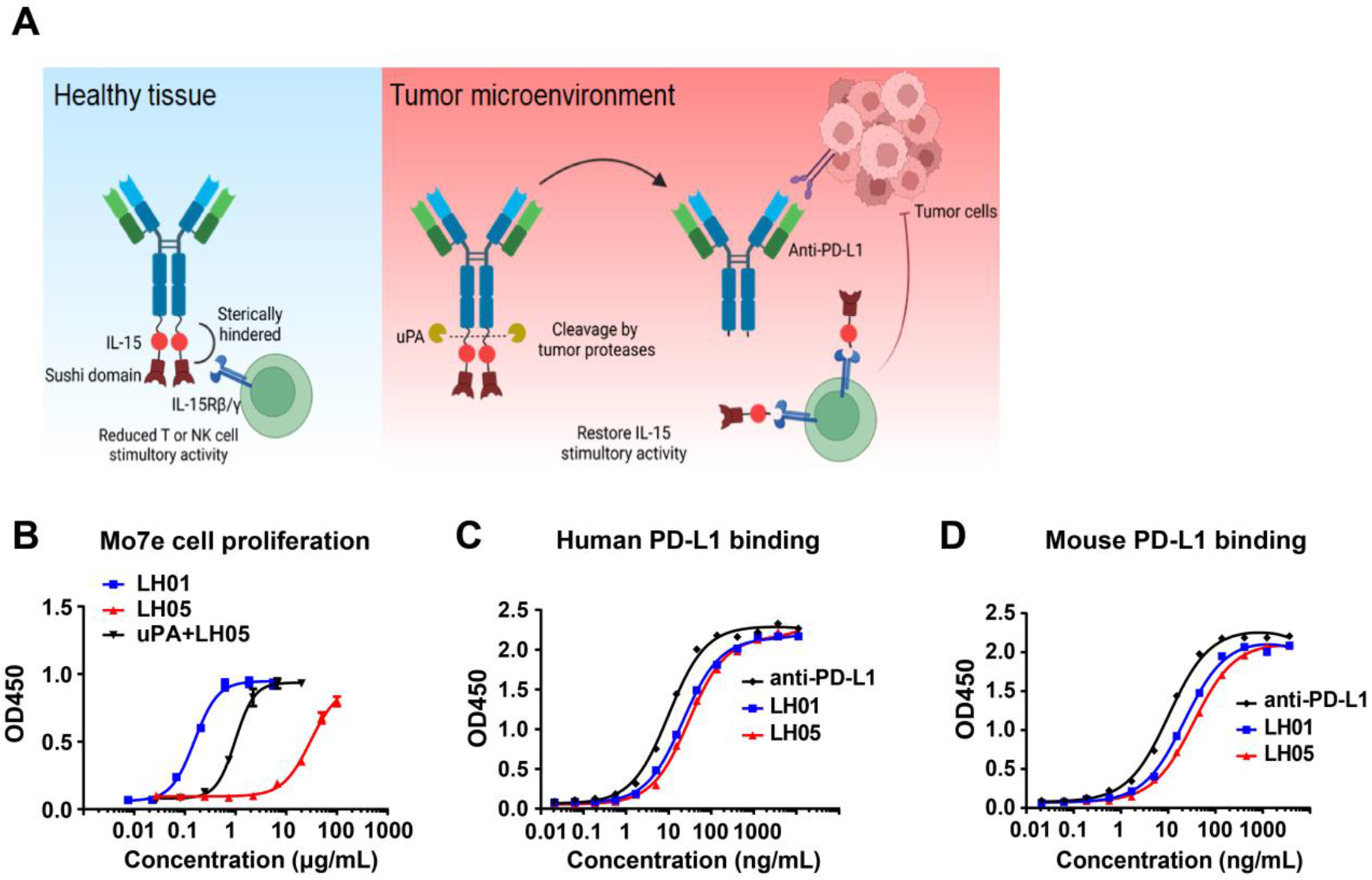
Structure based tumor-conditional anti-PD-L1/IL-15 design. **(A)** Schematic of the anti-PD-L1/IL-15 prodrug in healthy tissue and tumor environments. In healthy tissue, IL-15 is masked by the Fc fragment and sushi domain; in the tumor, it is cleaved by tumor-associated proteases, releasing the immunostimulatory ILR. **(B)** The proliferative potential of LH05 and uPA-cleaved LH05 (incubated with uPA at 20°C *in vitro* for 12 h) was compared with LH01 in human Mo7e cells. Data were analyzed using the four parameter fit logistic equation to calculate the EC_50_ values. **(C and D)** Binding of anti-PD-L1, LH01, and LH05 to plate-bound human (C) or mouse (D) PD-L1. Data were analyzed using the one site-total to calculate the EC_50_ values. All graphs are shown as mean ±SEM.

SDS-PAGE analysis revealed that LH05, but not LH03, can be cleaved after incubation with uPA (Supplementary Fig. 4C). LH05 (EC_50_ = 30.31 μg/mL or 147.5 nM) induced 168-fold less proliferative activity than LH01 (EC_50_ = 0.177 μg/mL or 0.88 nM) in Mo7e cells. When LH05 was cleaved, it restored the Mo7e cell proliferation stimulatory activity by more than 30 folds (EC_50_ = 4.9 nM) (Fig. 2B). LH05 also showed a decreased IL-2Rβ binding affinity than LH01 due to IL-15 masking, which could explain its weaker proliferative activity in human Mo7e cells (Supplementary Fig. 1). In ELISAs, both fusion proteins bound to human PD-L1 with a profile similar to that of the anti-PD-L1 antibody (EC_50_ = 21.83, 29.95, and 10.02 ng/mL, or 109.04, 145.80, and 69.28 pM, for LH01, LH05, and anti-PD-L1, respectively) (Fig. 2C), as well as similar affinities for mouse PD-L1 as anti-PD-L1 antibody (EC_50_ = 22.23, 34.90 and 10.32 ng/mL, or 111.03, 169.90 and 71.36 pM, for LH01, LH05, and anti-PD-L1, respectively) (Fig. 2D). Our results demonstrated that the anti-PD-L1 portion of LH05 was unaffected, and the ILR portion would be preferentially released within the TME to restore IL-15 activity.

### LH05 exhibits excellent safety profile in vivo

Given its lower immunostimulatory activity *in vitro*, we suppose that LH05 would attenuate the expansive capacity of peripheral lymphocytes and minimize systemic toxicity *in vivo*. To confirm whether LH05 has a significantly improved safety profile as compared with LH01, we treated mice with PBS, LH01 (5 mg/kg), or LH05 (10 mg/kg). After two LH01 treatments, all mice experienced dramatic body weight loss and eventually died within 6 days. Contrarily, none of mice treated with LH05 lost weight or died even after six injections (Fig. 3A and B). Compared with the PBS treatment, LH01 treatments induced a 229.3% increase in spleen weight, whereas double doses of LH05 only resulted in a 70.9% increase, indicating that LH05 can effectively shield IL-15 activity in circulation (Fig. 3C).

**Fig. 3.**
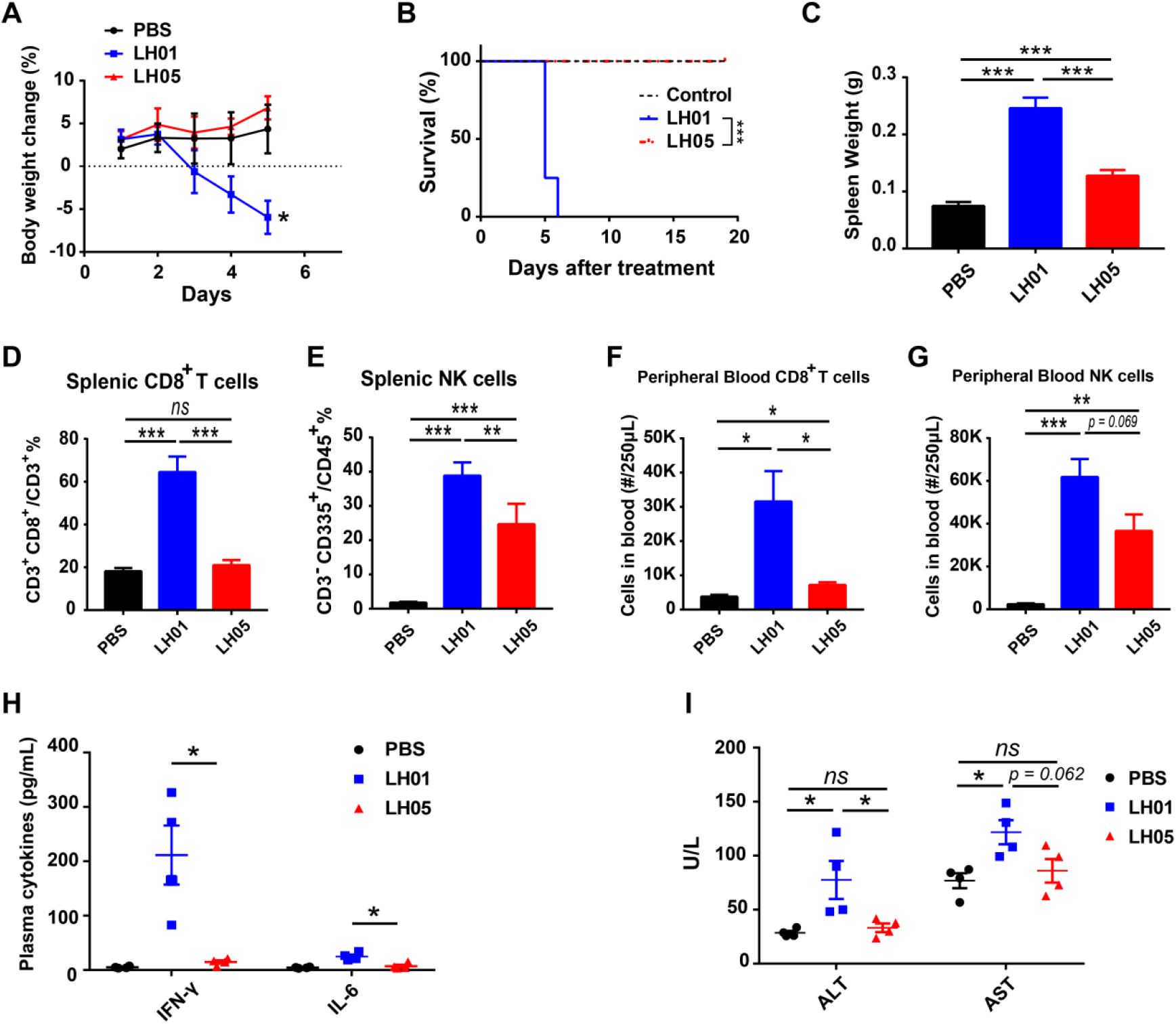
LH05 significantly reduces systemic toxicity. **(A and B)** Female Balb/c mice were intraperitoneally injected with PBS, LH01 (5 mg/kg), or LH05 (10 mg/kg) every 3 days, with body weight changes (A) and survival (B) monitored (n = 8). **(C)** Spleens of mice were extracted and weighed after euthanasia on day 5 (n = 4). **(D and E)** The percentages of splenic CD8^+^ T cells and NK cells are shown for populations of CD3^+^ and CD45^+^ lymphocytes, respectively. **(F and G)** The number of CD8^+^ T cells (F) and NK cells (G) in peripheral blood was counted. **(H and I)** Blood samples were collected after euthanasia on day 5, and plasma cytokine levels were measured using ELISA (H), ALT and AST plasma levels were also quantified (I). All graphs show the mean ±SEM. \**p* < 0.05; \*\**p* < 0.01; \*\*\**p* < 0.001; ns, not significant.

Interestingly, LH05 treatment did not lead to a significant increase in splenic CD8^+^ T proportion or peripheral blood CD8^+^ T-cell counts as compared to LH01 treatment (Fig. 3D-G, Supplementary Fig. 5). It retained some stimulatory activities on splenic and peripheral blood NK cells, although they were much weaker than those of LH01 (Fig. 3D-G, Supplementary Fig. 5). Moreover, unlike LH01, LH05 did not significantly trigger cytokines, such as IFN-γ and IL-6, further indicating that the risk of systemic toxicity induced by LH05 was greatly minimized (Fig. 3H). LH01 treatments also caused increased plasma alanine aminotransferase (ALT) and aspartate aminotransferase (AST) levels, whereas LH05 treatments did not cause any observed liver damage (Fig. 3I). Additionally, neither LH01 nor LH05 increased the plasma creatinine levels in comparison to PBS, implying that no renal injury occurred (Supplementary Fig. 6). Overall, these findings suggest that LH05 was effectively sheltered against peripheral activity and adverse effects.

### LH05 extends half-life due to the attenuated “cytokine sink” effect

In fact, even though being conjugated to a targeting moiety, wild-type immunocytokines may rapidly disappear from circulation before reaching tumor tissues due to the ubiquitous expression of their cognate receptors (known as the cytokine sink effect) [8]. Considering the significantly reduced affinity of the prodrug LH05 for the IL-15 receptor, it would confer superior pharmacokinetic properties to LH01. As expected, the plasma concentrations of LH05 decreased at a slower rate than those of LH01, with calculated half-lives of 8.40 and 3.45 h, respectively, following intravenous injection. These findings suggest that the reduced “cytokine sink effect” of the masked prodrug could prolong the half-life of LH01 by approximately 2.4 folds (Fig. 4A). To further investigate the tissue distribution of LH01 and LH05, mouse tissues were collected 18 h after treatment. The LH05 concentration in tumor tissue was significantly higher than the LH01 concentration, indicating its superior tumor-targeting capacity (Fig. 4B).

**Fig. 4.**
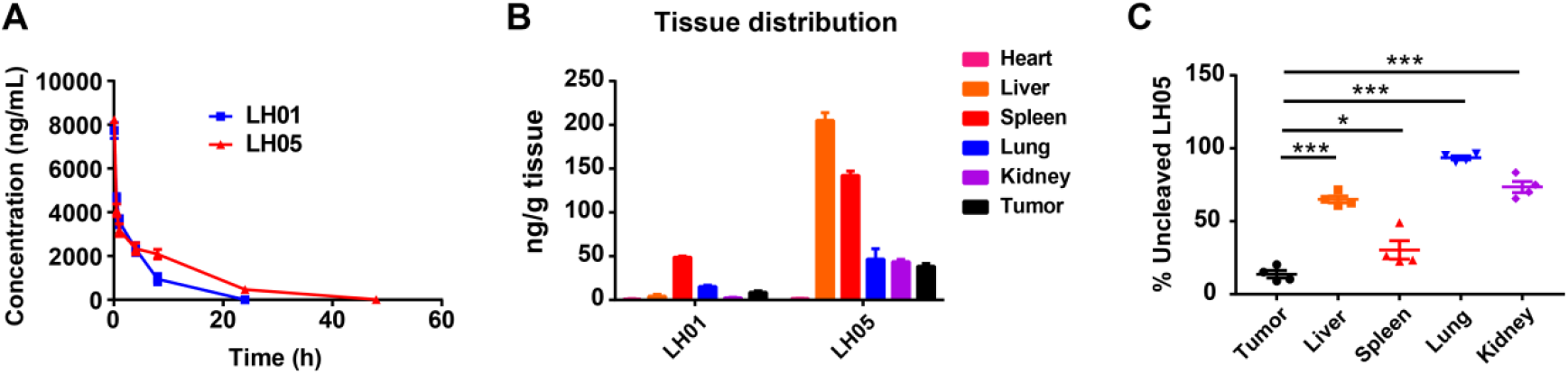
Prolonged half-life and improved tumor-targeting distribution of LH05. **(A)** Nine-week old male Balb/c mice were injected intravenously with 1 mg/kg LH01 or LH05 (equimolar molecules). The plasma concentration-time curves were plotted (n = 5). **(B)** RM-1 tumor-bearing mice (n = 4) were intravenously injected with 1 mg/kg LH01 or LH05, and tissues were collected at 18 h post-injection. The concentrations of LH01 or LH05 were measured using ELISA. **(C)** The cleavage efficiency of LH05 in tumor and organs were evaluated by determining the percentage of intact (un-cleaved) molecules in all detected anti-PD-L1 portions, using lysates collected after homogenization and centrifugation (n = 4). Both graphs show the mean ±SEM. \**p* < 0.05; \*\**p* < 0.01; \*\*\**p* < 0.001; ns, not significant.

Although uPA is reported to be highly expressed in multiple tumors, it is also found in normal tissues, such as liver, spleen, and kidney [24]. This poses a risk of non-selective cleavage of LH05 in healthy tissues, potentially limiting its therapeutic efficacy. Therefore, we further investigated the selectivity of LH05 cleavage between tumor and normal tissues, utilizing ELISAs coated with PD-L1 or IL-15Rβ to quantify total or un-cleaved LH05, respectively. Our results showed that LH05 was predominantly cleaved in tumor tissues when compared to any other tissue. Notably, we detected a relatively higher degree of cleavage of LH05 in the spleen, which was considerably lower than that observed in the tumor (Fig. 4C). These findings demonstrate the preferential cleavage of LH05 in tumors, reducing the risk of systemic toxicity.

### LH05 exhibits potent antitumor activity with greatly reduced toxicity

We first investigated the antitumor effects of LH05 in the syngeneic murine RM-1 prostate carcinoma model with a high uPA expression [25, 26]. LH01 was well tolerated at 2.5 mg/kg but it exerted much weaker antitumor activity than LH05. LH05 at 10 mg/kg was well tolerated and no mice had obvious weight loss. The LH02 (an IL-15 superagonist) dosage used in this study was 0.25 mg/kg, as established in previous research [22]. It is worth noting that LH05 also demonstrated superior antitumor efficacy as compared with anti-PD-L1+LH02 (Fig. 5A-C).

**Fig. 5.**
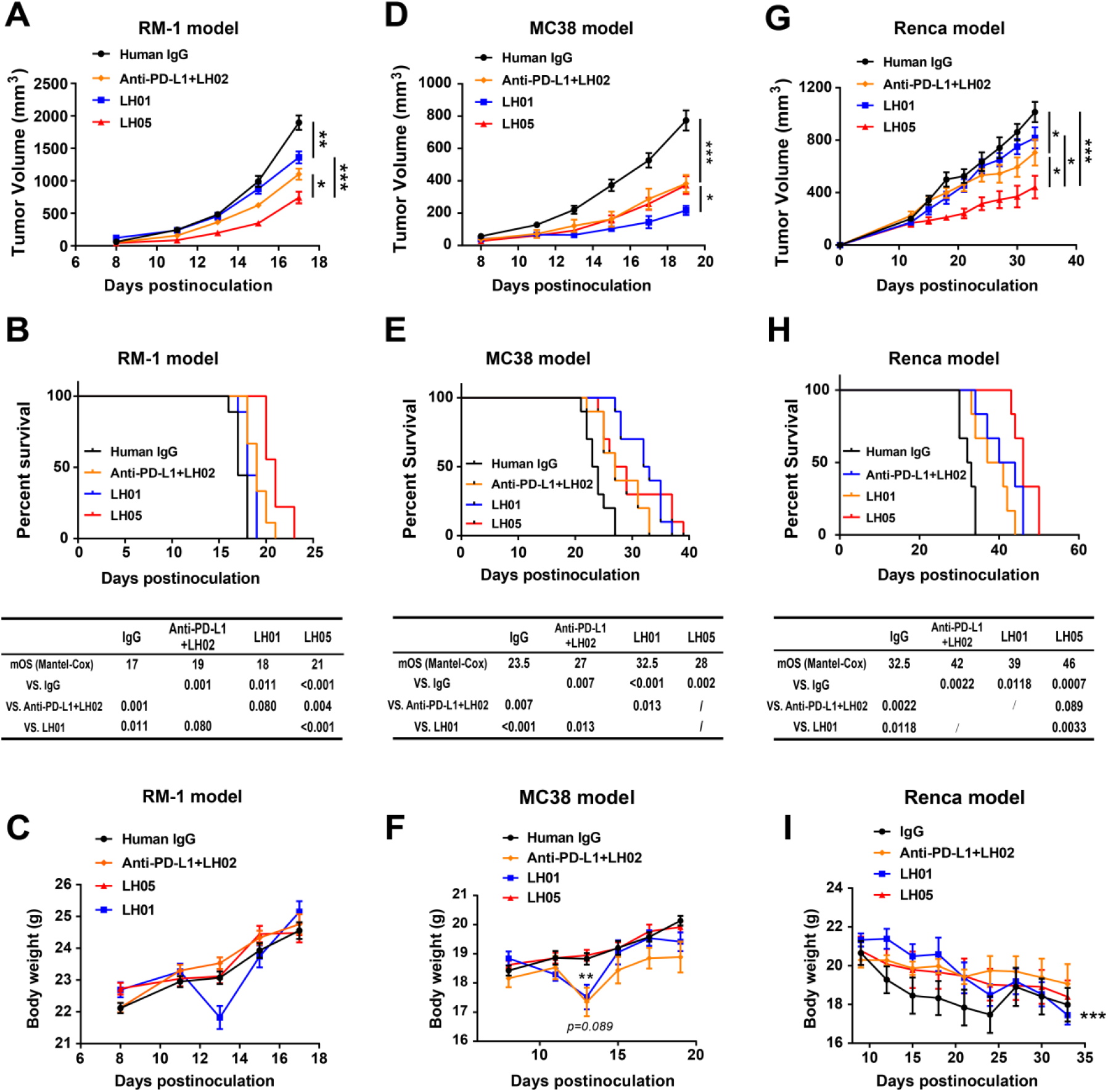
LH05 exerts potent antitumor efficacy with reduced toxicity. Mice were randomized into four groups based on tumor size, with treatment initiating when tumors reached 50-100 mm^3^. Tumor growth curves were plotted over time. Mice were observed for survival starting from the day after tumor cell inoculation. The body weights of tumor-bearing mice were recorded throughout the study. **(A-C)** RM-1 tumor cells (5 × 10^5^) were subcutaneously implanted into the right flank of male C57BL/6J mice. On days 9, 12, and 15 (n = 12), mice were intravenously injected with IgG control (10 mg/kg), anti-PD-L1 (10 mg/kg) + LH02 (0.25 mg/kg), LH01 (2.5 mg/kg) or LH05 (10 mg/kg). **(D-F)** MC38 tumor cells (5 × 10^5^) were subcutaneously implanted into the right flank of female C57BL/6J mice. On days 8, 11, 14, and 17 (n = 10), mice were intravenously injected with IgG control (10 mg/kg), anti-PD-L1 (10 mg/kg) + LH02 (0.25 mg/kg), LH01 (2.5 mg/kg) or LH05 (10 mg/kg). **(G-I)** Renca cells (5 ×10^5^) were suspended in 50 μL PBS and an equal volume of matrigel and subcutaneously implanted into the right flank of female Balb/c mice. On days 9, 12, 16, and 21 (n = 6), mice were intravenously injected with IgG control (10 mg/kg), anti-PD-L1 (10 mg/kg) + LH02 (0.25 mg/kg), LH01 (2 mg/kg) or LH05 (10 mg/kg). All graphs show the mean ± SEM. \**p* < 0.05; \*\**p* < 0.01; \*\*\**p* < 0.001; ns, not significant.

We further explored the antitumor effects of LH05 in the murine MC38 colon carcinoma model with a low uPA expression [16, 27]. LH05 exhibited comparable antitumor efficacy with anti-PD-L1+LH02 but it was somewhat weaker than LH01 (Fig. 5D). Although LH05 did not improve the median overall survival (mOS) as much as LH01 (28 vs 32.5), the difference was not statistically significant (Fig. 5E). Notably, LH01 and anti-PD-L1+LH02 induced significant body weight loss after two treatments, but not LH05, suggesting that the superior antitumor effects of LH01 was at the expense of severe toxicity (Fig. 5F). Moreover, in the murine Renca renal cell carcinoma model with a uPA expression level between MC38 and RM-1, LH05 generated a greater antitumor effect than LH01 or anti-PD-L1+LH02 (Fig. 5G and H) [27]. Besides, among the treatments, LH01 induced the most significant decreases in body weight in Renca tumor-bearing mice (18.14%, day 33 vs day 9) (Fig. 5I).

In all three tumor models, we observed that LH03 containing a non-cleavable linker displayed substantially weaker antitumor effects than LH05, demonstrating that *in vivo* cleavage is required to release the masked LH05’s bioactivity (Fig. 4A, D, and G, Fig. 1F, and Supplementary Fig. 2). These findings suggest that, compared with LH01, LH05 has superior tolerability while maintaining an uncompromised overall therapeutic effect in a proteolytic cleavage-dependent manner.

### LH05 induces both innate and adaptive immune responses for tumor control

In RM-1 tumor-bearing mice, anti-PD-L1+LH02 significantly increased the spleen weight as compared with IgG, but there was no obvious spleen weight gain observed in the LH05 treatment at a dose of 10 mg/kg (equivalent to 2.5 mg/kg of LH02 for IL-15), implying that LH05 had very weak peripheral immunostimulatory activity (Fig. 6A). A flow cytometry analysis of dissociated spleens and tumors from RM-1 tumor-bearing mice was then performed to explore the changes in splenic and intratumoral CD8^+^ T or NK populations. The gating strategy for the analysis of T and NK cells is shown in Supplementary Figs. 5 and 7. We observed that LH01, LH05, and anti-PD-L1+LH02 treatments markedly decreased the frequency of splenic CD4^+^ T cells as compared with the IgG treatment (Fig. 6B). The percentage of splenic CD8^+^ T cells increased significantly in the LH05 and anti-PD-L1+LH02 groups but not in the LH01 group (Fig. 6C). Furthermore, both LH05 and anti-PD-L1+LH02 treatments led to significantly decreased splenic CD4/CD8 ratio as compared to the other three treatments, indicating a stronger immune response (Fig. 6D). All other treatments markedly increased the splenic NK cells when compared to IgG treatment, but the percentage of NK cells was much lower in the LH01 group than in the LH05 group (Fig. 6E). Interestingly, although LH05 treatment significantly increased the percentage of splenic CD8^+^ T and NK cells, it did not induce spleen weight gain as compared to IgG treatment.

**Fig. 6.**
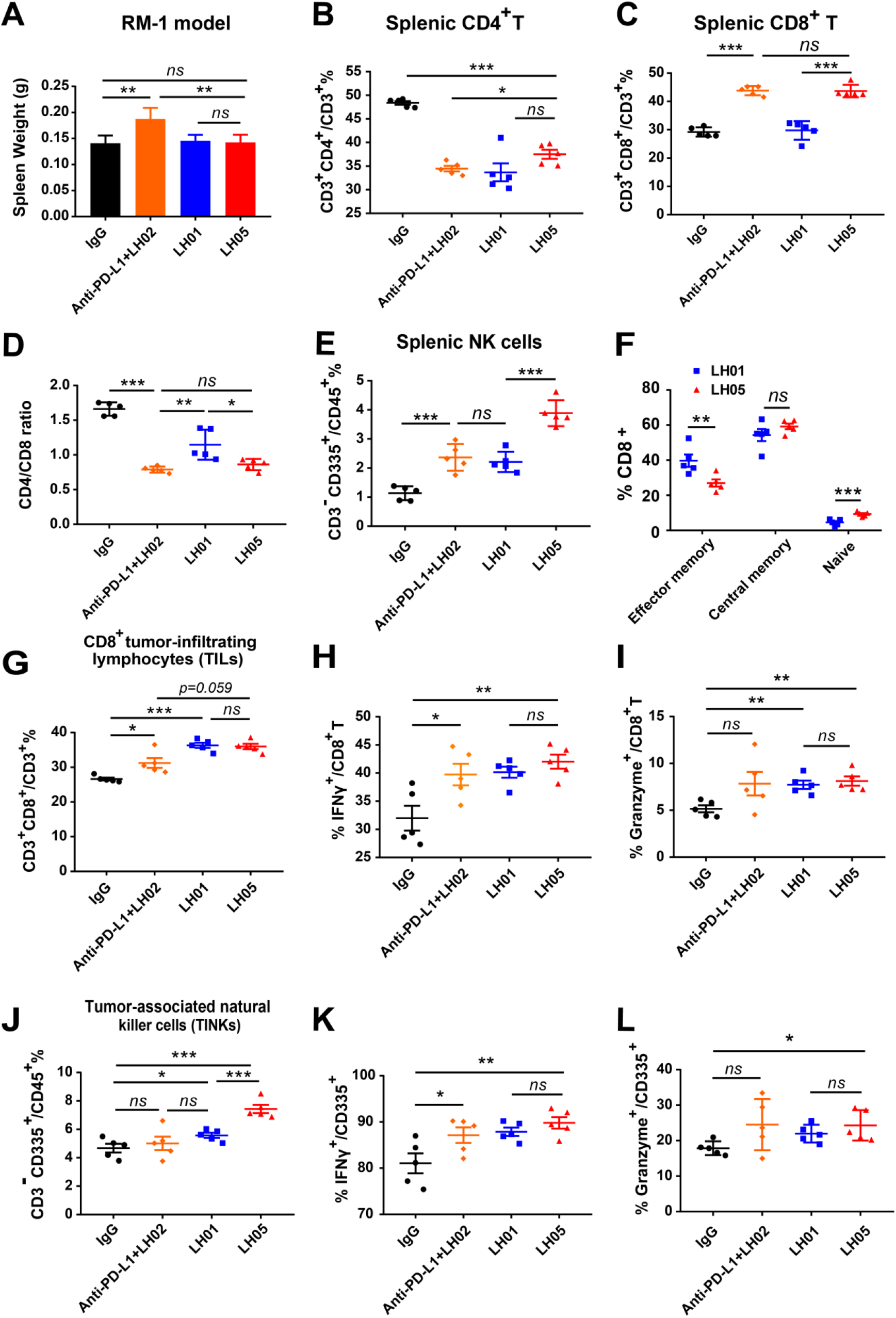
LH05 induces both adaptive and innate immune cells activation. Flow cytometry analysis of spleens and tumors of RM-1 tumor-bearing mice treated as described in Fig. 5. **(A)** The spleens of RM-1 tumor-bearing mice were extracted and weighed after euthanasia (n = 5). **(B and C)** The frequency of splenic CD4^+^ T cells (B) and CD8^+^ T cells (C) for CD3^+^ lymphocytes, respectively. **(D)** The ratio of CD4^+^ to CD8^+^ T cells was calculated. (**E**) The percentage of splenic NK cells for CD45^+^ lymphocytes was determined. **(F)** The expression of the memory cell markers CD62L and CD44 on splenic CD8^+^ T cells were assessed. **(G-I)** The percentage of intratumoral CD8^+^ T cells (G) within the population of CD3^+^ lymphocytes, and the frequency of IFNγ^+^ (H) or perforin^+^ (I) CD8^+^ T cells within the tumor were assessed. **(J-L)** The percentage of intratumoral NK cells (J) within the population of CD45^+^ lymphocytes, and the frequency of IFNγ^+^ (K) or perforin^+^ (L) NK cells within the tumor were determined. All graphs show the mean ±SEM. \**p* < 0.05; \*\**p* < 0.01; \*\*\**p* < 0.001; ns, not significant.

**Fig. 7.**
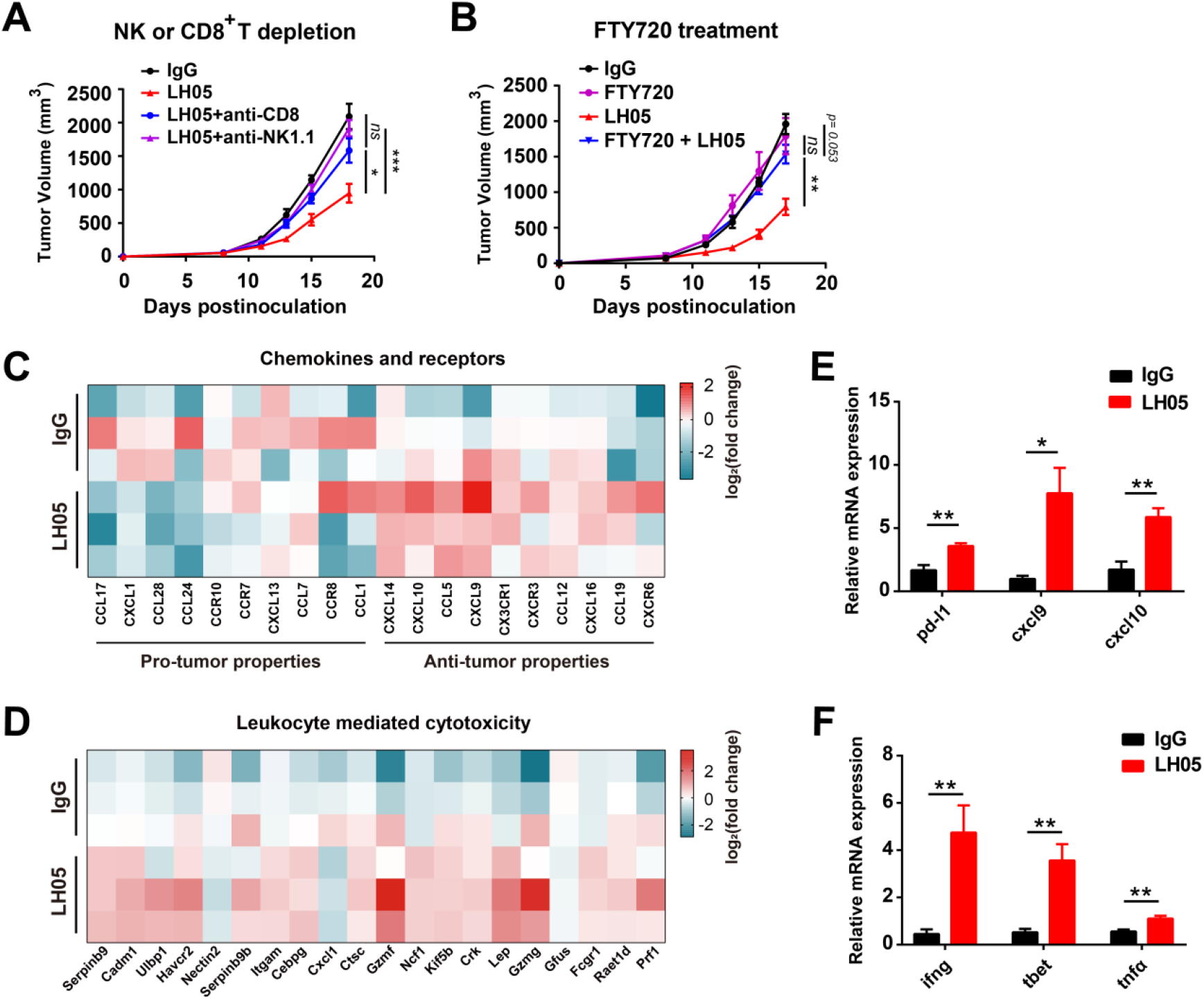
The recruited CD8^+^ T cells and NK cells contribute to LH05-mediated antitumor efficacy. **(A and B)** Growth curves of RM-1 tumors of mice treated with IgG, LH05, and CD8^+^ T or NK cells depletion (using anti-CD8 or anti-NK1.1 antibody, respectively) (A), or FTY720 (B) in the presence or absence of LH05. **(C and D)** The heatmaps depict gene expression alterations of chemokines and receptors (C) and leukocyte-mediated cytotoxic effector (D) in response to treatment, as indicated by the log2(fold change) values. (**E and F**) The expression levels of *pd-l1*, *cxcl-9*, and *cxcl-10* (E), and *ifng*, *t-bet*, and *tnfα* (F) in the TME were measured using quantitative real-time PCR. All graphs show the mean ± SEM. \**p* < 0.05; \*\**p* < 0.01; \*\*\**p* < 0.001; ns, not significant.

To further compare the difference in peripheral immunostimulatory activity between LH01 and LH05, we investigated the phenotypes of CD8^+^ T cells in the spleen. We observed that LH05 and LH01 treatments induced comparable proportions of central memory CD8^+^ T in the spleen, whereas LH01 treatment resulted in a significantly higher percentage of effector memory CD8^+^ T in the spleen as compared to LH05 treatment. Moreover, LH01 treatment markedly reduced the percentage of splenic naïve CD8^+^ T-cell population as compared to LH05 treatment, partly explaining why LH05 showed a better safety profile than LH01 (Fig. 6F).

LH05 treatment resulted in comparable increases in CD8^+^ tumor-infiltrating lymphocytes (TILs) than LH01 (Fig. 6G). To assess whether LH05 treatment enhanced the effector function of CD8^+^ T cells, we determined the IFN-γ and granzyme expression of tumor-infiltrating CD8^+^ T cells by flow cytometry. We found that LH05 treatment significantly increased the frequencies of both CD8^+^ IFNγ^+^ and CD8^+^ granzyme^+^ T cells inside the tumor as compared with IgG treatment, but no significant difference was observed between the LH05 and LH01 groups (Fig. 6H and I). However, LH05 treatment led to significantly higher levels of tumor-associated NK cells (TINKs) than LH01 treatment, which could explain why LH05 exhibited superior antitumor efficacy to LH01 (Fig. 6J). Similarly, LH05 treatment increased the frequencies of IFN-γ and granzyme-expressing CD335^+^ NK cells as compared to IgG treatment, but no significant difference was observed between the LH01 and LH05 groups (Fig. 6K and L). Altogether, these results showed that LH05 can be enzymatically cleaved within the TME, and the released ILR can activate the CD8^+^ T and NK cells for tumor inhibition.

### Both CD8^+^ T and NK cells recruited by LH05 contribute to its antitumor efficacy

First, we calculated the correlation coefficients of IL-15 expression and immune infiltration levels by employing the MCP-counter, xCELL, and CIBERSORT abs. mode algorithms. Then, we depicted the landscape of IL-15 correlating with immune cell infiltrates in various TCGA cohorts. Our resulting heatmap showed a statistically significantly positive correlation between IL-15 expression and immune infiltration of NK cells, particularly activated NK cells, and the central and effector memory subset of CD8^+^ T cells in the majority of cancers (Supplementary Fig. 8).

To ascertain which cell type contributes to LH05-mediated tumor control, we depleted the CD8^+^ T or NK cells in RM-1 tumor-bearing mice with respective depletion antibodies. The results showed that the depletion of NK cells completely abrogated the antitumor efficacy of LH05, indicating that NK cells played an essential role in tumor killing (Fig. 7A). Depleting CD8^+^ T cells also compromised LH05’s therapeutic effect, suggesting that CD8^+^ T cells are required for tumor immunity (Fig. 7A). We then used FTY720 (an inhibitor of T and NK cells egress from lymph nodes) to further determine whether the pre-existing immune cells within the tumor or recruited cells are indispensable for LH05’s anticancer effect. The experiment revealed that inhibiting lymph node egress almost entirely eliminated LH05’s efficacy (Fig. 7B). Additionally, the RM-1 tumor is known as a typical “cold tumor,” with few pre-existing T cells. Altogether, these findings suggest that LH05’s antitumor activity is primarily dependent on CD8^+^ T and NK cells that infiltrate the TME from the circulation, making it a promising candidate in the treatment of “cold tumors.”

To further evaluate the impact of LH05 treatment on immune responses, we conducted RNA sequencing (RNA-seq) of RM-1 tumors treated with or without LH05. A gene set enrichment analysis revealed that the LH05 treatment positively impacted the expression profile of chemokines and receptors in RM-1 tumors. Specifically, we observed a decrease in the expression of chemokines and receptors known for pro-tumor effects, such as CXCL1 and CCL28, whereas the expression of chemokines and receptors with antitumor properties, such as CXCL9 and CXCL10, was increased (Fig. 7C). These findings suggest that LH05 may potentially modulate the TME by altering the chemokine and receptor signaling balance toward an antitumor immune response. Additionally, we also discovered that LH05 treatment led to a general enhancement of immune pathways in the tumor tissue and an increase in the expression of genes related to leukocyte-mediated cytotoxic effector activity (Fig. 7D and Supplementary Fig. 9). These results are consistent with the findings from those obtained by flow cytometry (Fig. 6), indicating that LH05 have immunostimulatory effects on the TME.

We previously reported that the anti-PD-L1 treatment increased the *pd-l1* expression in tumors [22]. In the present study, the *pd-l1* level was significantly up-regulated by LH05, implying an improvement in antitumor immune responses. Two CXCR3 ligands, CXCL9 and CXCL10, are critical factors that facilitate immune cell migration to the TME and bring “heat” to tumors [28]. LH05 treatment resulted in a dramatic increase in *cxcl9* and *cxcl10* expression, which may explain the recruitment of CD8^+^ T cells in RM-1 tumors (Fig. 7E). Compared to IgG treatment, LH05 treatment also significantly increased the expression of *ifng*, *tnfα*, and *tbet* expression in the tumor, suggesting a T helper (Th) 1-skewed TME (Fig. 7F). IL-15 promotes intratumoral immune cell functions via a cytokine network involving XCL1, IFN-γ, CXCL9, and CXCL10 [29]. Taken together, when LH05 reaches the TME, the reactivated LH05, specifically the released ILR, can stimulate an immune-activating microenvironment by recruiting CD8^+^ T and NK cells, promoting their expansion and cytotoxicity, and inducing Th-1 type cytokines secretion to exert a potent antitumor immunity.

### LH05 restores response to immunotherapy in U251 cold tumors

Glioblastoma (GBM) is a highly malignant primary brain tumors with a five-year survival rate of <5% despite treatment by surgical resection, targeted radiation therapy, and chemotherapy [30]. GBMs are considered “cold” tumors characterized by poor lymphocyte infiltration and an immunosuppressive TME, which poses challenges for ICIs to stimulate effective antitumor immune responses [31].

Given LH05’s ability to overcome ICI resistance and re-induce immunotherapeutic responses, we evaluated its antitumor efficacy in the U251 glioblastoma xenograft model. LH01 was used as a control with a dosage of 3 mg/kg for safety concerns. The results showed that LH01 exhibited only a minor antitumor effect without statistically significant differences when compared to that of PBS, indicating that it failed to elicit a robust immune response to inhibit tumor growth. Contrarily, the LH05 treatment significantly reduced the tumor volume and weight compared with PBS or LH03 (10 mg/kg) controls, indicating that LH05 cleaved in the TME can trigger profound antitumor immunity and efficiently suppress tumor growth (Fig. 8A and B). At 10 mg/kg, LH05 was well tolerated and did not cause weight loss in the U251 tumor-bearing mice (Supplementary Fig. 10). Furthermore, LH05 treatment significantly reduced Ki67 expression of tumors when compared with the other three treatments, demonstrating a reduced tumor cell proliferation and metastasis ability (Fig. 8C). Overall, LH05’s ability to overcome immunotherapy resistance and stimulate antitumor immunity in the glioblastoma model highlights its potential as a therapeutic strategy for other tumors types with similar immunosuppressive characteristics.

**Fig. 8.**
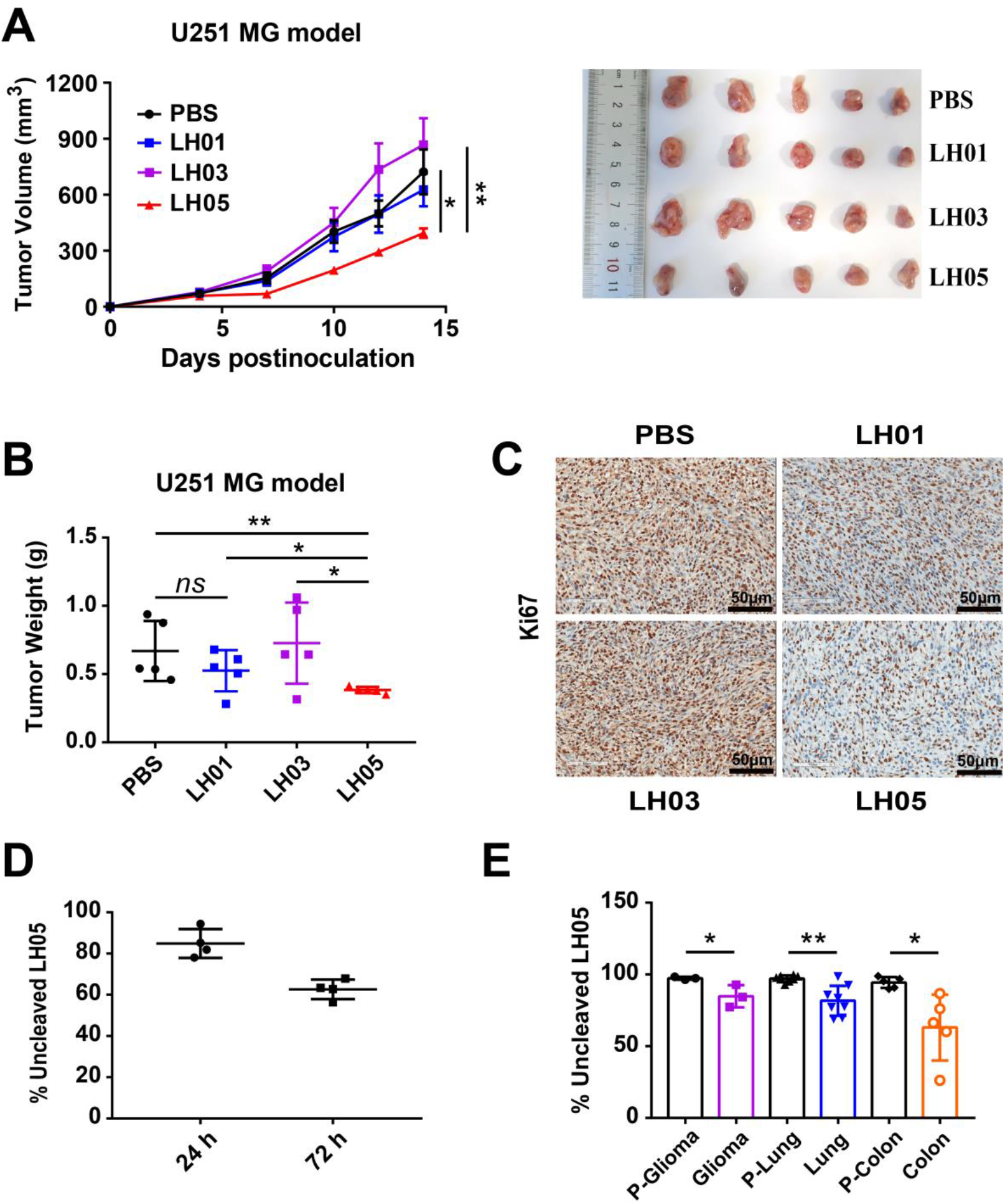
LH05 exerts enhanced antitumor efficacy than LH01 in U251 cold tumor. **(A)** NCG mice were inoculated subcutaneously with 2 × 10^6^ U251 cells and received 4.0 × 10^6^ fresh human PBMCs intravenously on day 4. Mice were then randomized into four groups, and treatment initiated when tumors reached 50-100 mm^3^. The groups were treated with PBS, LH03 (10 mg/kg), LH01 (3 mg/kg), or LH05 (10 mg/kg) intraperitoneally on days 5, 8, and 11 (n = 5). Tumor volumes were measured. **(B)** After euthanasia, tumors were removed, weighed, and photographed. **(C)** Immunohistochemical staining for Ki67 was performed on the tumor tissues to assess cell proliferation. **(D)** LH05 was incubated with human serum at 37°C for 24 or 72 hours before the cleavage was measured by ELISA (n = 4). **(E)** LH05 was incubated with human cancer homogenate or adjacent normal tissues homogenate at 37°C for 24 hours, and the cleavage efficiency was detected by ELISA. Data are shown as the mean ± SEM. \**p* < 0.05; \*\**p* < 0.01; \*\*\**p* < 0.001; ns, not significant.

### LH05 is stable in human serum and susceptible to tumor-specific proteolytic cleavage

Given the marked increase in protease expression in solid human tumors, we verified LH05’s efficient and selective cleavage in a range of primary human tumor samples. We first incubated LH05 with human serum from healthy donors (n = 4) for 24 or 72 hours. LH05 underwent slight cleavage after incubation for 24 hours, with only an approximately 40% cleavage rate observed after 72 hours (Fig. 8D), demonstrating a good human serum stability.

To evaluate the specificity of LH05 cleavage by tumors versus healthy tissues, we obtained various tumors and their corresponding peri-tumoral tissues from patients. We homogenized these tissues and incubated the homogenates with LH05; then, ELISA was performed to quantify the un-cleaved and total LH05. As expected, the cleavage efficiency rates varied across individuals. Some colonic tumors efficiently cleaved LH05 (>40% within 24 hours), but the lung or glioma tumors exhibited only 35% or even a lower cleavage after 24 hours of digestion (Fig. 8E). Notably, no LH05 cleavage was observed in any of the adjacent normal tissue homogenates, indicating that LH05 is stable in human normal tissues (Fig. 8E). Overall, these data suggest that LH05 has a low risk of systemic toxicity due to its high peripheral stability and can be specifically activated in human tumors. However, careful consideration of tumor types is crucial to guarantee efficient cleavage *in vivo*.

## Discussion

Immunocytokines are designed to enhance the targeting activity of cytokines, but only a modest 10-fold increase in targeted activity is reportedly achieved, which provides a limited increase in the therapeutic index [32]. In clinical studies, the majority of immunocytokines still has a dose-limiting toxicity similar to the parental cytokines [33]. To achieve more effective modalities of immunocytokines, further reducing systemic toxicity and increasing antitumor activity are imperative.

One solution for reducing systemic toxicity of immunocytokines is to engineer cytokines with reduced affinity for their cognate receptors [34–36]. Decreasing affinity toward the cognate receptor can reduce the “cytokine sink” effect, thereby extending half-life. Additionally, the lower biological activity allows for higher doses and immunocytokine accumulation at the tumor site. For example, IL-2 has been engineered to reduce its affinity for IL-2Rα or IL-2Rβ/γ [37–39]. However, these mutants reduced the affinity of immunocytokines for both tumoral and peripheral lymphocytes, posing a challenge to the balance between insufficient antitumor activity at low doses and the risk of systemic toxicity at high doses.

Prodrug-based strategies for conditionally activating cytokines in the TME can potentially improve their safety profile while maintaining the antitumor activity. One of the most promising directions for achieving tumor-localized cytokine activation is by leveraging tumor-associated proteases. Until now, two main prodrug strategies have been developed for macromolecules, including monoclonal antibodies, using a masking domain or via steric hindrance. Various masking domains have been used to shield cytokines, including native cytokine receptors, antibody fragments, anti-cytokine antibodies, and peptides [12]. Fu et al. have reported cognate receptor-masked IL-2, IL-12, IL-15, and IFN-α prodrugs [13–16]. WTX-124, an IL-2 prodrug, comprising native human IL-2 linked to a Fab antibody fragment (inactivation domain) and a single-domain antibody targeting human albumin (half-life extension domain), has entered phase I clinical trial (NCT05479812) by Werewolf [40]. However, the released masking moiety might still bind to the activated cytokine due to its high affinity, and the introduction of the masking domain could complicate the structure and increase the immunogenicity risk.

Currently, there is limited research on immunocytokine prodrugs. Only Askgene has reported an anti-PD-1/IL-15 prodrug, ASKG915, that utilizes IL-2Rβ to mask the IL-15 activity [41]. In this study, we propose a next-generation immunocytokine prodrug strategy with two features: 1) a novel steric hindrance method is used to mask cytokine activity; and 2) the cytokine would not be confined to the antibody moiety but be released after a tumor-associated proteolysis. With this strategy we constructed a tumor-conditional anti-PD-L1/IL-15 immunocytokine, LH05, which has a prolonged plasma half-life and improved safety profile due to the attenuated “cytokine sink” effect in circulation. As expected, it exhibited a potent antitumor efficacy in a proteolytic cleavage-dependent manner with significantly lower systemic toxicity than wild-type anti-PD-L1/IL-15. Our results showed that our design has the following clear advantages: it does not introduce additional proteins or peptides, thereby avoiding an increase in structural complexity or the potential for immunocytokine immunogenicity; and after cleavage, the released ILR can elicit broad-spectrum immune responses with superior antitumor efficacy.

Mechanically, the excellent efficacy of LH05 can be attributed to both the PD-L1-trans delivery of ILR to the TME and the release of active ILR after cleavage. Previously reported PD-1 cis-targeted IL-2/IL-15R agonists, including PD-1-laIL-2, αPD1-IL15m, and αPD1-IL15-R, can selectively deliver IL-2 or IL-15 to PD-1^+^CD8^+^ TILs and bypass NK cells [42–44]. All of these immunocytokines showed an antitumor efficacy that was dependent on intra-tumoral CD8^+^ T cells but not on NK or lymph node T cells. However, in our study, the IL-15 superagonist ILR was trans-delivered into the TME by the anti-PD-L1 moiety, and it not only stimulated the adaptive and innate immune cells but also increased their infiltration into tumor tissues, illustrating a more comprehensive antitumor role than the PD-1-cis-delivered immunocytokines.

RM-1 prostate carcinoma and U251 glioblastoma are both considered immunologically “cold” tumors, where therapeutic difficulties and failures are primarily due to an immune-hostile and immunosuppressive TME that abrogates T-cell infiltration and activation. LH05 showed significant antitumor effects in both of these models, indicating its potential as a treatment for cold tumors. To further explore why LH05 is effective, we investigated the TME. CXCL9 and CXCL10 are two critical chemokines for recruiting effector T cells from the circulation into the tumor and establishing a “hot” TME [27]. Our findings revealed that LH05 treatment significantly increased the mRNA levels of CXCL9 and CXCL10 but it had no influence on the Treg recruiting CCL-17 and CCL-22 (Supplementary Fig. 11) [45]. Additionally, there was a significant increase in the IFNγ, TNF-α, and T-bet expression levels after LH05 treatment, suggesting a Th1-biased TME. Moreover, the greatly improved safety of conditionally activated LH05 allows the use of higher doses, which is beneficial for improving antitumor effects.

In conclusion, LH05 represents a new class of next-generation immunocytokines that differ from all the reported molecules, including conditionally activated cytokines or immunocytokines [12]. In preclinical models, it demonstrated a favorable safety profile and superior antitumor efficacy. LH05 has a great potential for creating a “hot” TME by recruiting lymphocytes from the circulation into the tumor and inducing a Th1-skewed TME. Therefore, it is a promising candidate for further clinical investigation in patients with ICIs resistance or cold tumors. However, individual differences in tumor-associated protease expression levels, including uPA, MMPs, or matriptase, could add uncertainty to the clinical application of such products, which should also be considered in all prodrug strategies.

## Materials and Methods

### Cloning, expression, and purification

LH01, LH02, and anti-PD-L1 were constructed and produced as previously described[22]. For LH03 or LH05 construction, the human IL-15 mutant (IL-15N72D)/IL-15Rα-sushi domain (Ile31 to Val 115) complex (ILR) was fused to the C-terminal of anti-PD-L1 heavy chain via a GS flexible linker or a uPA-substrate linker, respectively, in the pMF09 vector we reported before [46]. The light and heavy chain expression plasmids of LH03 or LH05 were mixed with 25 kDa linear polyethylenimine (PEI, Polysciences) and transiently transfected in HEK293E cells. All fusion proteins were purified by using a protein A affinity column (GE Healthcare) and analyzed on SDS-PAGE in the reducing condition.

### Cell lines

HEK293E and Mo7e cell lines were kept in our laboratory and cultured as previous described [22]. RM-1 murine prostate carcinoma cell line, MC38 murine colon carcinoma cell line, Renca murine renal carcinoma cell line, and U251 human glioblastoma cell line were obtained from the American Type Culture Collection (ATCC). RM-1, MC38, Renca, and U251 cells were maintained in Dulbecco’s modified Eagle’s medium (DMEM) containing 10% FBS. All of the cells mentioned above were kept in aseptic conditions and incubated at 37°C with 5% CO_2_.

### ELISA to evaluate the affinity of anti-PD-L1/IL-15 for PD-L1

ELISAs were conducted following standard procedures. Briefly, 96-well ELISA plates (Corning) were coated with 1.0 μg/mL of recombinant human or mouse PD-L1 (Novoprotein) overnight at 4°C, followed by washing four times with PBST (PBS, 0.05% Tween-20) and blocked with 5% bovine serum albumin for 2 hours at room temperature. After washing the plates, serial dilutions (1:3) of LH01, LH05, or anti-PD-L1 antibody were added in duplicate to the plates and incubated at room temperature for 2 hours. The plates were washed four times and then incubated with Peroxidase AffiniPure Goat Anti-Human IgG (H+L) (Jackson ImmunoResearch, 1:10,000 dilution) at room temperature for 1 hour. After washing, the plates were incubated with TMB single component substrate solution (Solarbio) in the dark for 3-5 min. The reaction was stopped with 2 M sulfuric acid, and absorbance was read at 450 nm with a reference at 630 nm (Teacan, Infinite 200 PRO).

### Pharmacokinetics studies of immunocytokines

Plasma samples were collected 5 min, 0.5 h, 1 h, 4 h, 8 h, 24 h, and 48 h after intraperitoneal injection with 1 mg/kg LH01 or LH05. A 96-well ELISA plate, previously coated overnight at 4°C with 1.0 μg/mL of recombinant human PD-L1, was incubated for 2 hours with plasma samples from mice. The following experimental procedure for ELISA was the same as described above.

### Quantitative biodistribution studies of immunocytokine

Heart, liver, spleen, lung, kidney, and tumor tissues of RM-1 tumor-bearing mice were collected. About 25 mg of tissues were weighed and homogenized in 10% PBS before being centrifuged to obtain supernatant. We employed two ELISA assays to quantify the amount of LH01 or LH05, either cleaved or un-cleaved, in each homogenate, normalized by total tissue weight. And the above ELISA assay developed to evaluate anti-PD-L1/IL-15 affinity for PD-L1 was used to detect the total amount of LH01 or LH05 in both cleaved and un-cleaved forms. Since both LH01 and LH05 can bind IL-15Rβ with high affinity, an ELISA assay was developed to detect LH01 or un-cleaved LH05 with IL-15Rβ coated on the plate and Peroxidase AffiniPure Goat Anti-Human IgG (H+L) (Jackson ImmunoResearch, 1:10,000 dilution) as the detection antibody. In detail, 2.0 μg/mL of recombinant human IL-15Rβ was coated overnight at 4°C, and then incubated with supernatant from tissue homogenate for 2 hours. The following experimental procedure for ELISA was the same as described above.

### *In vitro* cleavage of immunocytokines with uPA

*In vitro* cleavage was performed by incubating 10 μg LH03 or LH05 with 0.25 μg uPA in phosphate buffer saline in a total reaction volume of 10 μL at 20°C for 12 h.

### LH05 stability in human serum and cleavage of LH05 by human tumors

Human serum was purchased from Shanghai Xinfan Biotechnology Co., Ltd. 1 μL LH05 (2 μg) was added to 9 μL human serum, then incubated at 37 °C for 24 or 72 hours. Colon, lung and brain tumors as well as their adjacent peritumoral tissues were collected from Zhejiang University School of Medicine, Shanghai Jiao Tong University School of Medicine, and Fudan University School of Medicine, respectively. All subjects provided broad informed consent for the research use of their biological samples. Homogenization was performed using FastPrep tissue homogenizer (MP Bio, USA). Supernatant was collected by centrifugation at 10000 g for 15 min. For the cleavage experiments, 9 μL tissue lysate was incubated with 0.2 μg LH05 (0.2 mg/mL) at 37°C for 24h.

The un-cleaved and total LH05 was quantified by the ELISA described above. To confirm the feasibility of the above ELISA methods, LH03 containing non-cleavable linker and uPA-activated LH05 were included as negative and positive control, respectively.

### Mo7e cell proliferation assay

Mo7e cells were washed with human GM-CSF free medium (RPMI1640 + 10% FBS) before being seeded into 96-well plates at a density of 2×10^4^ cells in a volume of 50 μL per well. After 4 hours’ starvation, serial dilutions (1:3) of LH01, LH03, or LH05 (treated with or without uPA) were added to the plate in sextuplicate at 50μL per well to achieve a final density of 2×10^4^ cells/100 μL/well. Cell viability was measured using the Cell Counting Kit-8 kit (Dojindo, Japan) after 96 hours of incubation at 37°C with 5% CO_2_. The absorbance was read at 450 nm with an ELISA reader (Teacan, Infinite 200 PRO), and the final OD450 value of the sample wells have subtracted the blank reading.

### Animal experiments

All animal experiments were approved by the Animal Care and Use Committee of Shanghai Jiao Tong University. Sex-matched Balb/c and C57BL/6 mice aged 6-8 weeks were purchased from Shanghai SLAC Laboratory Animal Co., Ltd. Female NCG mice aged 6-8 weeks were purchased from Jiangsu GemPharmatech LLC. All mice were raised in pathogen-free environments and received humane treatment throughout the experimental period. Human peripheral blood mononuclear cells (PBMCs) were purchased from Shanghai Milestone Biotechnologies. For antitumor studies, tumors were measured every two or three days using a digital caliper, and volumes were calculated as (length×width^2^)/2. Tumor Growth Inhibition (TGI): TGI (%) = 100 ×(1-T/C). T and C were the mean tumor volumes of the treated and control groups, respectively.

### Flow cytometry analysis

About 150 mg of tumor tissues was cut into small pieces and re-suspended in digestion buffer [RPMI1640 medium containing collagenase IV (2 mg/mL) and hyaluronidase (1.2 mg/mL)]. Tumors were digested for 60 min at 37°C and then filtered through a 200-mesh nylon net to obtain the cell suspension. The cells were washed by RPMI 1640 and filtrated through a 200-mesh nylon net again, and then resuspended in FACS buffer (PBS + 2% FBS) to obtain pre-treated single cell suspension. Splenic lymphocytes were isolated from the spleens with lymphocyte separation medium (Dakewe, China) after the spleens were gently ground.

Cell samples were blocked with anti-mouse CD16/CD32 mAb 2.4G2 (BD Biosciences, USA) at 4°C for 30 min before being incubated with surface marker antibodies at 4°C for 30 min. The Zombie Red Fixable Viability Kit (BioLegend) was used to exclude dead cells. For the detection of intracellular IFN-γ and granzyme, cell samples were further fixed and permeabilized by Fixation/Permeabilization Kit (BD Biosciences).

The following antibodies and reagents were used: mouse anti-mouse CD45.2-APC-Cy7 (BD Biosciences), hamster anti-mouse CD3e-FITC (BD Biosciences), rat anti-mouse CD4-PE (BD Biosciences), rat anti-mouse CD8a-BV510 (BD Biosciences), rat anti-mouse CD8a-APC (BD Biosciences), rat anti-mouse Nkp46-BV421 (BioLegend), rat anti-mouse Nkp46-Alexa Flour 647 (BD Biosciences), rat anti-mouse IFN-γ-BV786 (BD Biosciences), mouse anti-human/mouse Grazyme B-PE Cyanine 7 (BioLegend), rat anti-mouse CD8-FITC (BioLegend), rat anti-mouse CD44-PE (BioLegend), rat anti-mouse CD62L-APC (BD Biosciences). Flow cytometry was performed using CytoFLEX cytometer (Beckman Coulter, USA) or ACEA Novocyte (Agilent, Technologies, USA) and analyzed by FlowJo 10 (TreeStar, USA) or NovoExpress (Agilent, Technologies, USA).

### Detection of plasma ALT, AST, CREA, UREA, IFN-γ, and IL-6

Plasma levels of ALT, AST, CREA, and UREA were measured with a Roche biochemical analyzer (Roche, Switzerland). Plasma levels of IFN-γ and IL-6 was determined by mouse IFN-γ and IL-6 ELISA Kit (Multi Sciences, China) according to the manufacturer’s procedures, respectively.

### Depletion of immune cells in mice

To deplete the individual immune cell types, RM-1 tumor-bearing mice were intravenously injected with IgG control (10 mg/kg) or LH05 (10 mg/kg) on days 9, 12, and 15. For cell depletion, mice were intraperitoneally given 200 μg of anti-NK1.1 antibody (BioXcell, BE0036) or 200 μg anti-CD8α antibody (BioXcell, BE0061) on days 7, 9, and 13. Tumor growth curves (n = 5) were plotted. To study the effect of lymphocytes egressing from lymph nodes, RM-1 tumor-bearing mice were administered with IgG (10 mg/kg, i.v.), LH05 (10 mg/kg, i.v.), or FTY720 (Sigma-Aldrich, USA) with or without LH05. FTY720 (25 μg) was intraperitoneally administered every other day beginning 8 days after tumor cell inoculation (n = 6).

### RNA isolation and qRT-PCR analysis

Total RNAs were extracted from tissues using the Ultrapure RNA Kit (Cwbio, China), and cDNA was synthesized using a PrimeScript RT Master Mix (Takara, Japan). Real-time qRT-PCR was performed on an Applied Biosystems 7500 Fast Real-Time PCR System (Thermo Fisher Scientific, USA) using Hieff qPCR SYBR Green Master Mix (Yeasen, China). The primer sequences are listed in Table S1. All results were normalized to GAPDH expression and calculated using the 2^-(△△Ct)^ method.

### Immunohistochemistry analysis

The tumor tissues were fixed in 4% paraformaldehyde, embedded in paraffin, and sectioned (4 μm). After dewaxing and hydration, heat-induced epitope retrieval and 3% H_2_O_2_ treatment were used to block endogenous peroxidase activity. The tumor sections were then blocked with 5% BSA for 30 min before being incubated with anti-human Ki67 rabbit antibody (Servicebio, China) at 4°C overnight. Next, the sections were incubated with the HRP-conjugated goat anti-rabbit secondary antibody (Servicebio, China) for 60 min. Finally, the sections were stained with a DAB detection kit (Dako, Copenhagen, Denmark) and hematoxylin, then observed and photographed with an OLYMPUS BX53 Microscope.

### Statistical analysis

Prism 7.0 software (GraphPad, USA) was used for statistical analysis. The two-tailed Student’s t test and analysis of variance were used to determine the statistical significance of differences between experimental groups (*: p < 0.05, **: p < 0.01, ***: p < 0.001). The log rank (Mantel-Cox) test was used to assess survival.

## Competing interests

The authors declare no competing interests.

## Notes

### Competing Interest Statement

The authors have declared no competing interest.

