## Supplemental Figures for "Next-generation anti-PD-L1/IL-15 immunocytokine elicits superior antitumor immunity in cold tumors with minimal toxicity"

Wenqiang Shi *et al.*

Corresponding to:.

**Fig. S1-S11**

**Table S1**

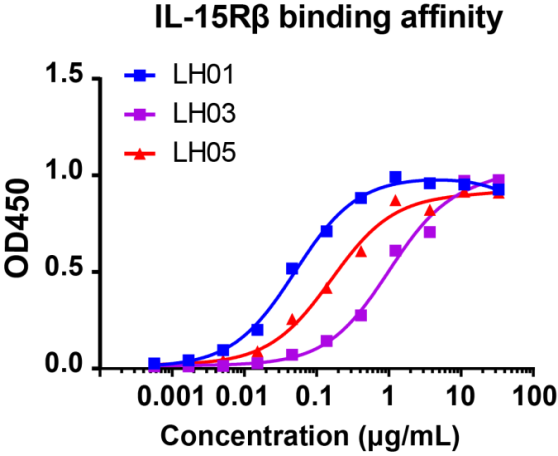

**Supplementary Figure 1.** Binding of LH01, LH03, and LH05 to plate-bound human IL-15Rβ. Data were analyzed using the one site-total to calculate the affinity values to be 0.21, 5.23, and 0.85 nM, respectively.

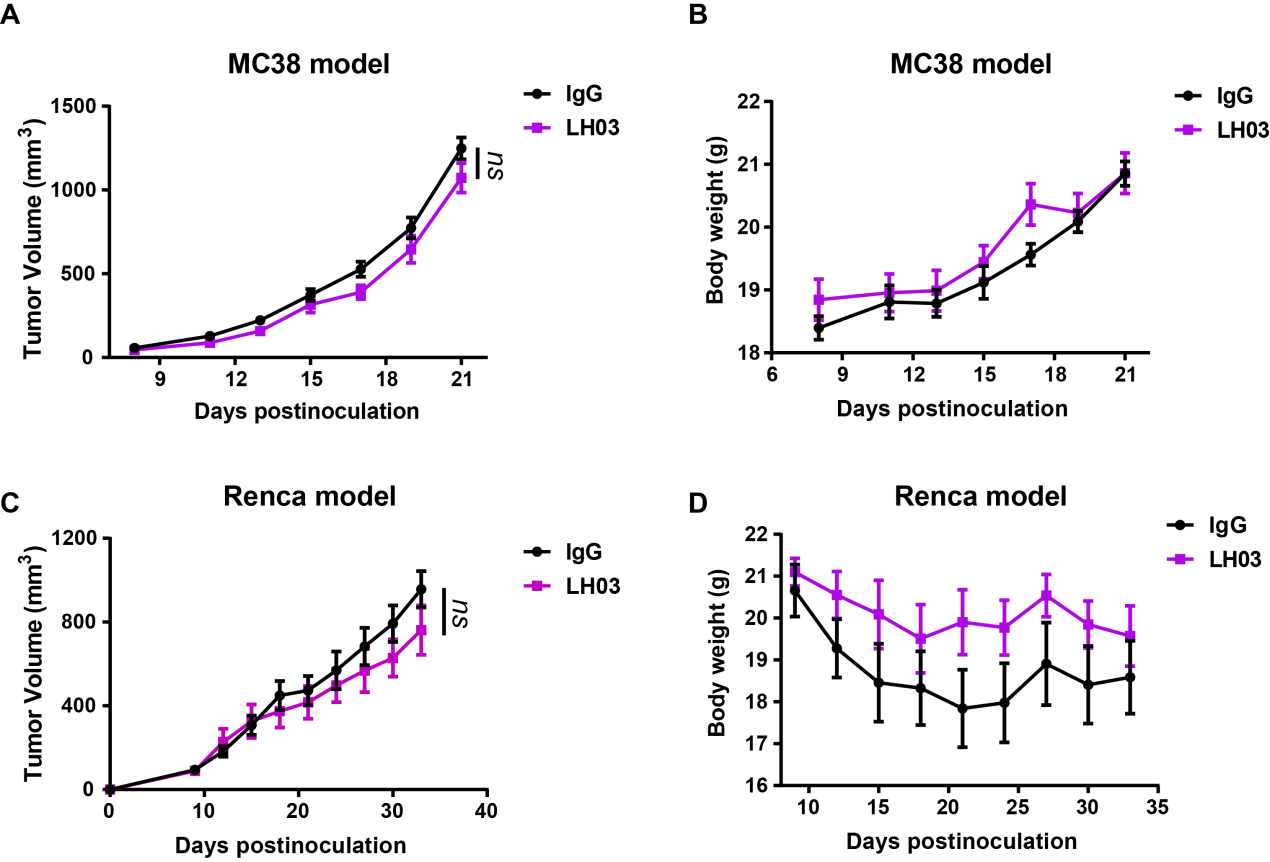

**Supplementary Figure 2. The engineered concealed IL-15-R demonstrates no significant antitumor effects despite its good safety profile.**

**(A-B)** MC38 tumor-bearing mice were intravenously injected with human IgG (10 mg/kg) or LH03 (10 mg/kg) on days 8, 11, 14, and 17 (n = 8). The tumor progression curves of tumor volumes were plotted (A) and body weights (B) were recorded. **(C-D)** Renca tumor-bearing mice were intravenously injected with human IgG (10 mg/kg) or LH03 (10 mg/kg) on days 9, 12, 16, and 21 (n = 7). The tumor progression curves of tumor volumes were plotted (C) and body weights (D) were recorded.

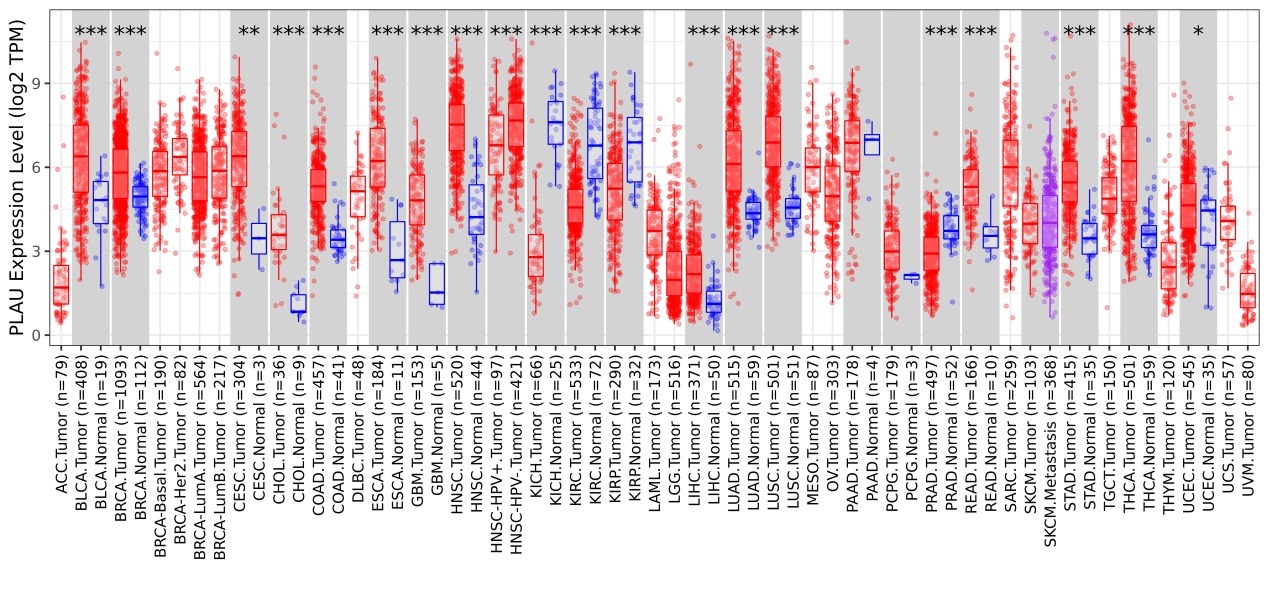

**Supplementary Figure 3.** **Human uPA expression level in tumors and adjacent normal tissues.** The DiffExp module of TIMER (Tumor IMmune Estimation Resource) website online analysis of the comparison of uPA expression levels for all samples from TCGA (The Cancer Genome Atlas). Red dots indicate tumor tissues, and blue dots indicate adjacent normal tissues.

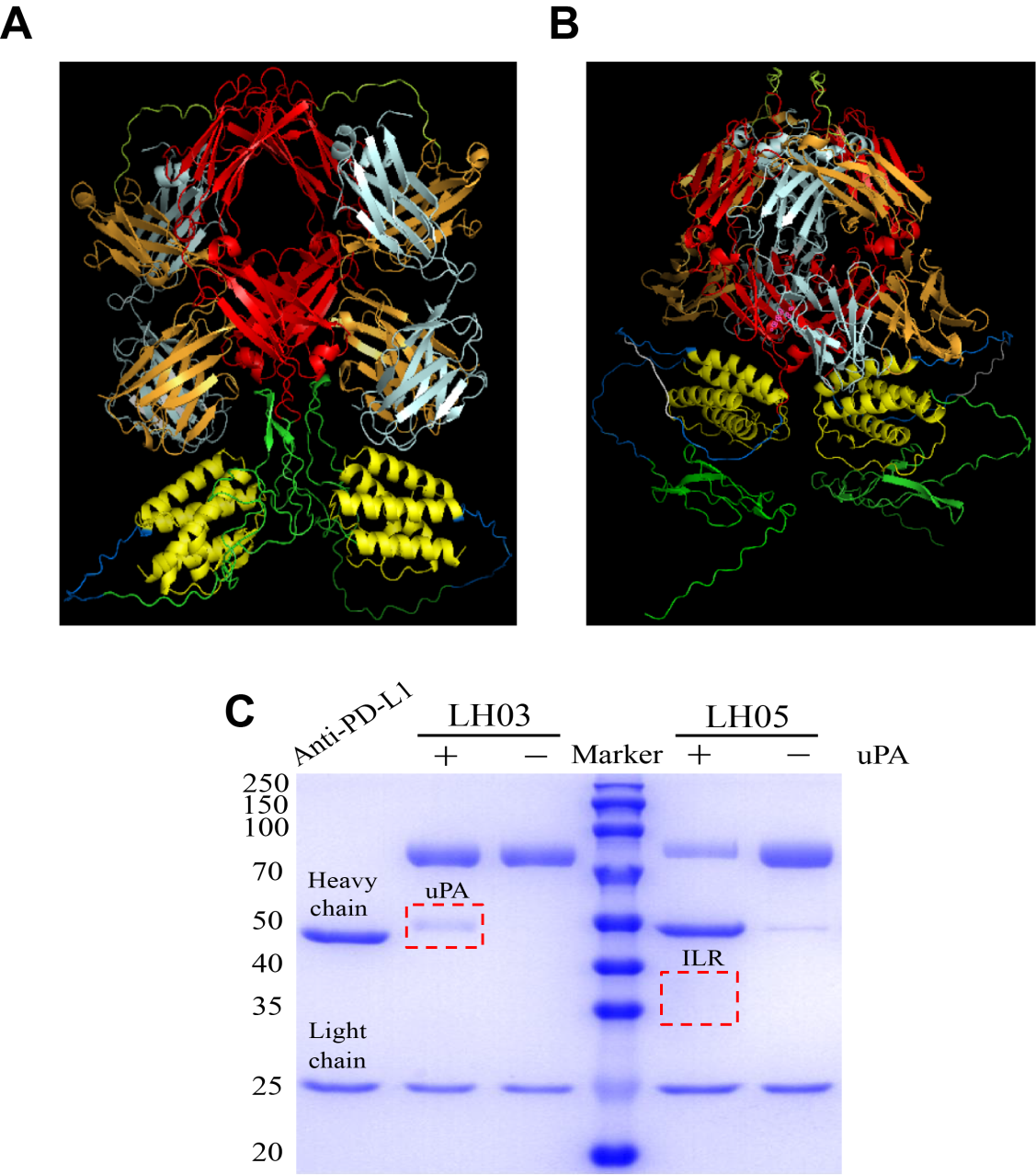

**Supplementary Figure 4. Engineering a tumor-conditional anti-PD-L1/IL-15. (A and B)** The structures of LH01 (A) and LH05 (B) were predicted using AlphaFold 2 v2.0, a machine learning-based computational method that can predict protein structures with reasonable accuracy. The light chains of anti-PD-L1 are indicated in pale cyan, the VH and CH1of anti-PD-L1 in orange, the Fc fragments of anti-PD-L1 in red, the IL-15Rα-sushi domain in green, IL-15 in yellow, GS linker in marine, and the protease-cleavable linker in white. The model provides a static representation of a plausible conformation of LH01 or LH05. **(C)** Reducing SDS-PAGE analysis of anti-PD-L1, LH03 or LH05 as well as uPA-cleaved LH03 or LH05. The experiment was repeated three times independently with similar results.

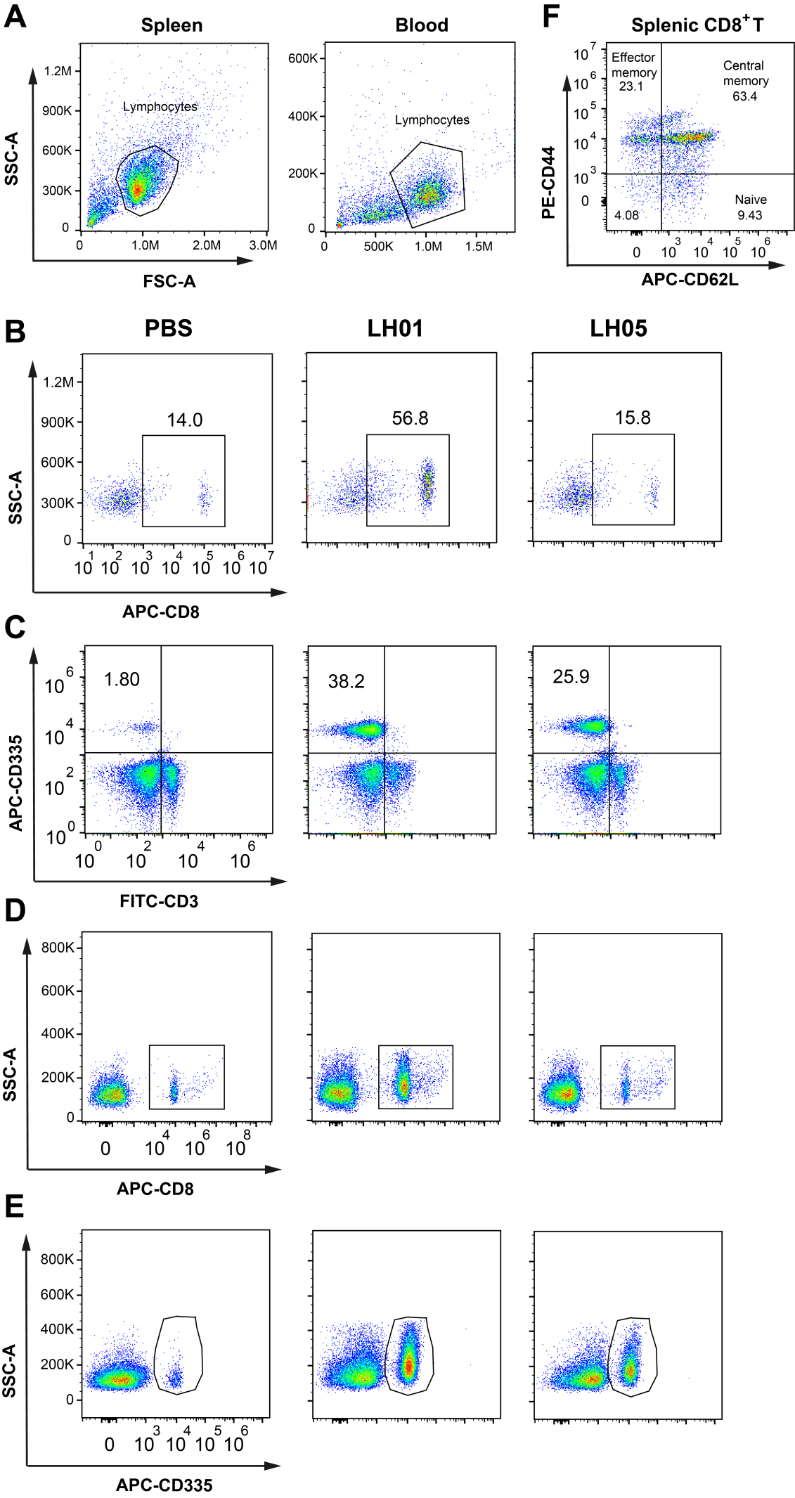

**Supplementary Figure 5.** Flow cytometric analysis. (A) The gate to identify lymphocytes of spleen or peripheral blood. (B-C) The percentages of splenic CD8^+^ T cells and NK cells for CD3^+^ or CD45^+^ lymphocytes were shown. (D-E) The cell counts of CD8^+^ T cells or NK cells in peripheral blood were shown. (F) The percentages of effector memory T cells were shown for populations of splenic CD8^+^ T cells. Data are shown as the mean ± SEM.

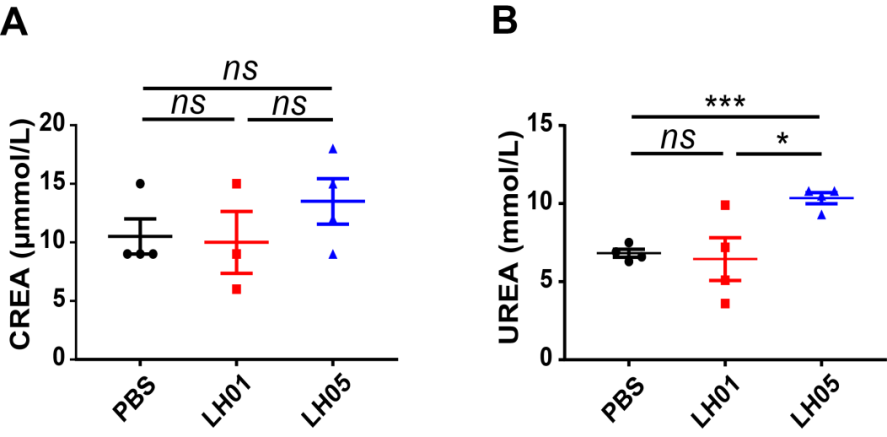

**Supplementary Figure 6.** The levels of CREA and UREA in the plasma were quantified. All graphs show the mean ± SEM.

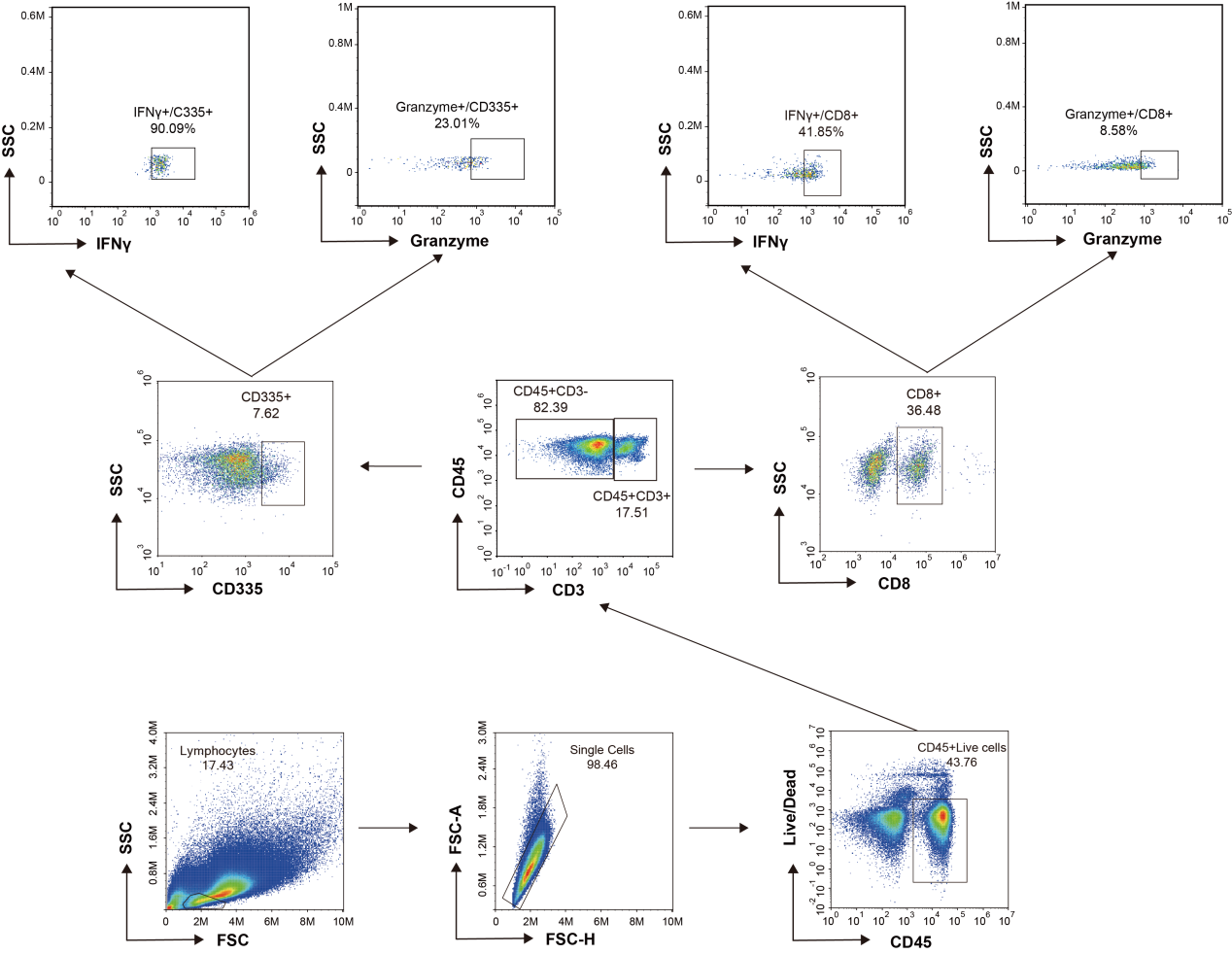

**Supplementary Figure 7.** Representative gating strategy for identifying CD8^+^ T cells and NK cells in the tumor tissue.

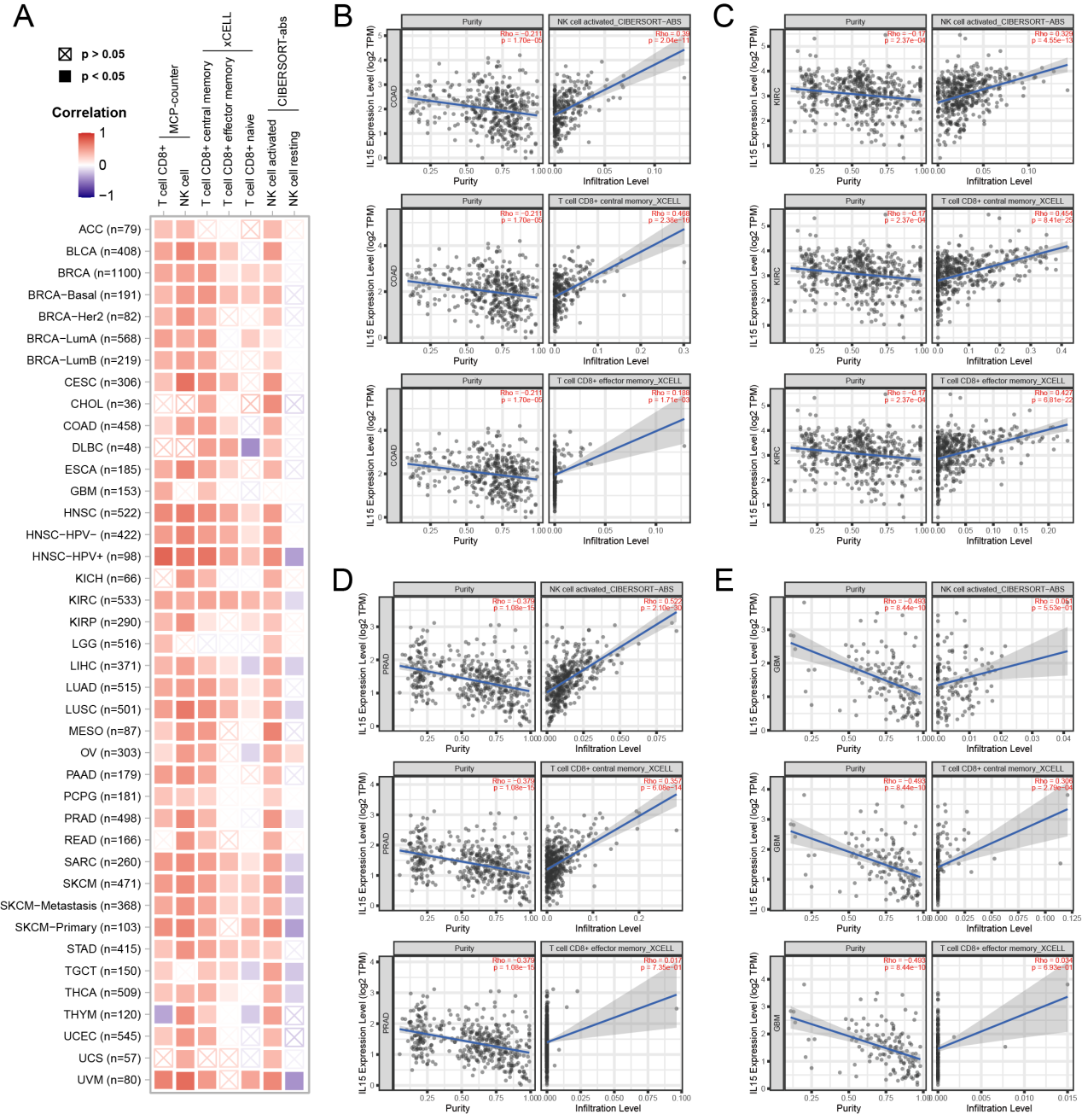

**Supplementary Figure 8.** IL-15 expression correlates with the immune cell infiltration in tumors. **(A)** Correlations between IL-15 expression level and the infiltration levels of CD8^+^ T and NK cells across TCGA cancers. Scatter plots of activated NK, central and effector memory subset of CD8^+^ T cells infiltration levels related to IL-15 expression in **(B)** COAD, **(C)** KIRC, **(D)** PRAD, and **(E)** GBM tumors were presented. TIMER2.0 database was utilized to explore the correlation between IL-15 levels and immune cell infiltration in all TCGA cancers.

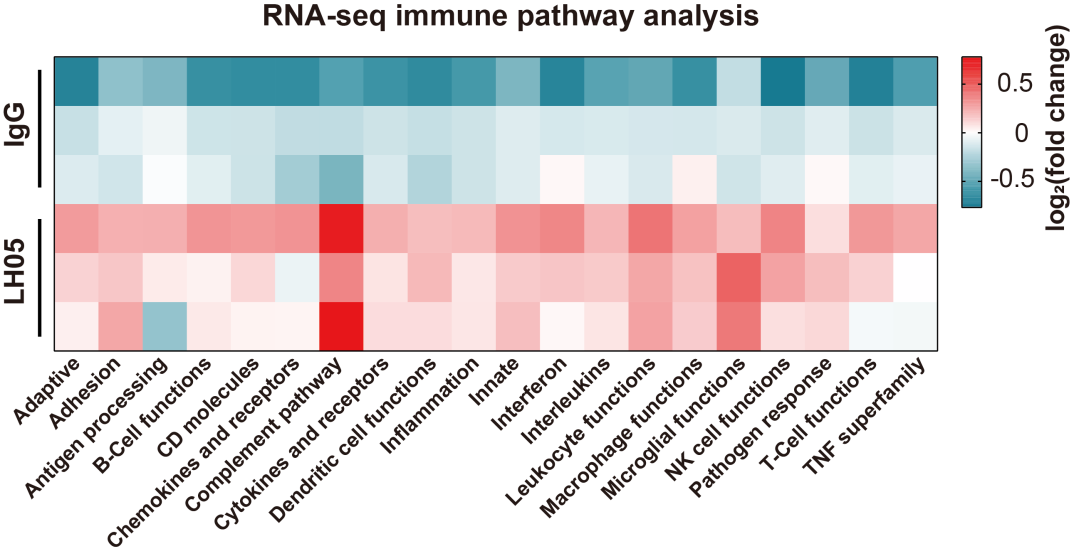

**Supplementary Figure 9.** Heatmap plot displays pathway scores of the 20 immune-related pathways.

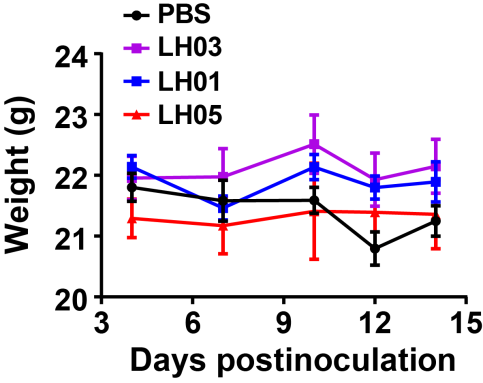

**Supplementary Figure 10.** Body weights of mice were monitored.

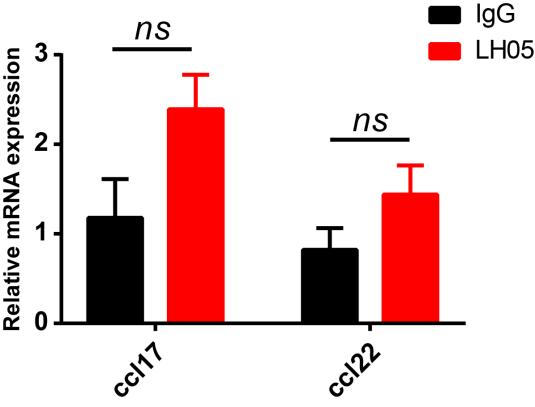

**Supplementary Figure 11.** The expression levels of *ccl-17* and *ccl-22* in the TME were measured using quantitative real-time PCR. All graphs show the mean ± SEM. **p* < 0.05; ***p* < 0.01; ****p* < 0.001; ns, not significant.

**Table S1.** Primer sequences for quantitative real-time PCR analysis

| Gene | Forward | Reverse |
| --- | --- | --- |
| *Pd-l1* | GCTCCAAAGGACTTGTACGTG | TGATCTGAAGGGCAGCATTTC |
| *Cxcl9* | GGAGTTCGAGGAACCCTAGTG | GGGATTTGTAGTGGATCGTGC |
| *Cxcl10* | CCAAGTGCTGCCGTCATTTTC | GGCTCGCAGGGATGATTTCAA |
| *Ccl17* | TACCATGAGGTCACTTCAGATGC | GCACTCTCGGCCTACATTGG |
| *Ccl22* | AGGTCCCTATGGTGCCAATGT | CGGCAGGATTTTGAGGTCCA |
| *Ifng* | ATGAACGCTACACACTGCATC | CCATCCTTTTGCCAGTTCCTC |
| *Tnfα* | CCCTCACACTCAGATCATCTTCT | GCTACGACGTGGGCTACAG |
| *Tbet* | AGCAAGGACGGCGAATGTT | GGGTGGACATATAAGCGGTTC |
| *Gapdh* | TCTCCTGCGACTTCAACA | TGGTCCAGGGTTTCTTACT |
